# Signatures of Discriminative Copy Number Aberrations in 31 Cancer Subtypes

**DOI:** 10.1101/2020.12.18.423278

**Authors:** Bo Gao, Michael Baudis

## Abstract

Copy number aberrations (CNA) are one of the most important classes of genomic mutations related to oncogenetic effects. In the past three decades, a vast amount of CNA data has been generated by molecular-cytogenetic and genome sequencing based methods. While this data has been instrumental in the identification of cancer-related genes and promoted research into the relation between CNA and histo-pathologically defined cancer types, the heterogeneity of source data and derived CNV profiles pose great challenges for data integration and comparative analysis. Furthermore, a majority of existing studies have been focused on the association of CNA to pre-selected “driver” genes with limited application to rare drivers and other genomic elements.

In this study, we developed a bioinformatics pipeline to integrate a collection of 44,988 high-quality CNA profiles of high diversity. Using a hybrid model of neural networks and attention algorithm, we generated the CNA signatures of 31 cancer subtypes, depicting the uniqueness of their respective CNA landscapes. Finally, we constructed a multi-label classifier to identify the cancer type and the organ of origin from copy number profiling data. The investigation of the signatures suggested common patterns, not only of physiologically related cancer types but also of clinico-pathologically distant cancer types such as different cancers originating from the neural crest. Further experiments of classification models confirmed the effectiveness of the signatures in distinguishing different cancer types and demonstrated their potential in tumor classification.

## Introduction

Copy number variations (CNV) are a class of structural genomic variants, in which the regional ploidy differs from the normal state of the corresponding chromosome. Germline copy number variations constitute a major part of genomic variability within and between populations and are an important contributor to genetic and inherited diseases [1]. In most cancer types, an extensive number of somatic CNV, usually referred to as sCNV or CNA (copy number aberrations), accumulate during the progression of the disease [2, 3]. CNAs have been shown to be directly associated with the expression of driver genes [4, 5] where expression of oncogenes can be increased by copy number amplifications and tumor suppressor genes can be suppressed through heterozygous or homozygous deletions. On a genomic level, recurrent patterns of CNVs are observed in a number of cancer types and have been associated with cancer prognosis and development [6].

While traditional karyotyping and various DNA hybridization techniques had provided insights into specific CNV events, the systematic, genome-wide screening for CNV emerged in the 1990s as a reverse in-situ hybridization technology termed “comparative genomic hybridization” (CGH; [7, 8]). Chromosomal CGH allowed the semi-quantitative profiling of copy number changes over complete (tumor)genomes. However, the technology was limited through its chromosomal banding based low resolution [9] and only indirect association of CNV events with putative target genes. The hybridization of tumor (or germline) DNA to genome-spanning matrices was refined through the use of substrates (”arrays”) containing thousands to more than 2 millions of mapped DNA sequence elements [10, 11], now allowing the direct association of the experimental read-out to specific genome features. A variation of this principle, SNP (single nucleotide polymorphisms) based arrays [12], originally developed for the detection of allelic variations within populations, was rapidly adopted for CNV detection and allelic decomposition analysis in cancer [13]. Nowadays, Next-Generation Sequencing (NGS) techniques are increasingly adopted to detect copy number variations, [14, 15, 16] although technologies with coverage below shallow whole genome sequencing [17] show reduced utility for the analysis of CNV events when compared to high-density arrays. Regardless of their technical heterogeneity, a large number of CNV data has been generated in the past three decades, which represents an invaluable asset for genomics studies.

Spurred by an increasing interest in genomic heterogeneity as well as mutational patterns shared across tumor types, the exponential growth of available cancer CNV profiling data stimulated the study of patterns in the context of meta-analyses and large consortia studies, across multiple groups of tumors [18, 6, 3, 19, 20, 21, 22, 23]. The majority of the studies focused on finding either associations to cancer driver genes or the impact of focal regions in specific tumor types. As a result, the CNA patterns were often characterized by the coverage of driver genes in contrast to comparative analyses of the whole genome. While this approach can provide direct connections to the established theories, it also bears two drawbacks. First, the distribution of cancer driver genes is extremely skewed: a few hallmark drivers are responsible for a large percentage of tumorigenesis, while a long-tail of rare or putative drivers are reckoned to be the cause of the rest [24]. Second, besides the association with somatic mutations, researches have also discovered different facets of CNA in their relations to cellular regulations and genome dynamics [25, 26, 27, 28]. Therefore, CNA patterns that solely rely on driver genes often lack the capacity to embrace the full spectrum of the aberrations. To generate CNA patterns, which capture their multitude of uniqueness, it would be more comprehensive to abstract by the aberrations’ characteristics, rather than using focal regions that overlap with driver genes.

In translational research, an imminent goal of studying the CNA patterns of different cancers is to gain insight in designing new therapeutic protocols. Recently, the discovery of circulating cell-free DNA (cfDNA), which presents a potential for early-stage and non-invasive cancer detection, attracts the attention of both the academia and the industry [29, 30, 31, 32]. While many cfDNA methods in cancer detection are focusing on somatic mutations, it is believed that the ultimate solution would come from an ensemble of different genomic aberrations [33]. One of such aberrations is CNA in cancers that bear high burdens of copy number mutation. Large-scale cancer genome studies have illustrated the recurrent CNA patterns across different types of cancer. Several studies have employed CNA of cfDNA as biomarkers and demonstrated their potential in identifying cancer types and tissues of origin [34, 35, 36]. As an emerging field, the accurate identification of genomic abnormalities and classifications of the cfDNA yet remains challenging, and the characterization of CNA patterns across cancer types would provide a valuable piece of the solution to the puzzle.

Due to the technical heterogeneity and underlying biological variability, the meta-analysis of multi-platform cancer CNV data, which is necessary for comprehensive “pan cancer” studies, also poses great challenges in data integration and normalization [37]. In this study, we have assembled a collection of 56,077 CNA profiles and created the CNA signatures of 31 cancer subtypes, where each signature represented the CNA landscape of a specific cancer group. The signatures were generated from a computational pipeline (Figure 1), which unifies heterogeneous data and is powered by a hybrid model of neural networks and attention algorithm. Using the signatures, we also constructed a multi-label classifier of cancer types and tissues of origin from copy number segmentation data. The result illustrates the genetic uniqueness of CNA in different cancer types and demonstrates the potential of CNA signatures in tumor identification.

**Figure 1:**
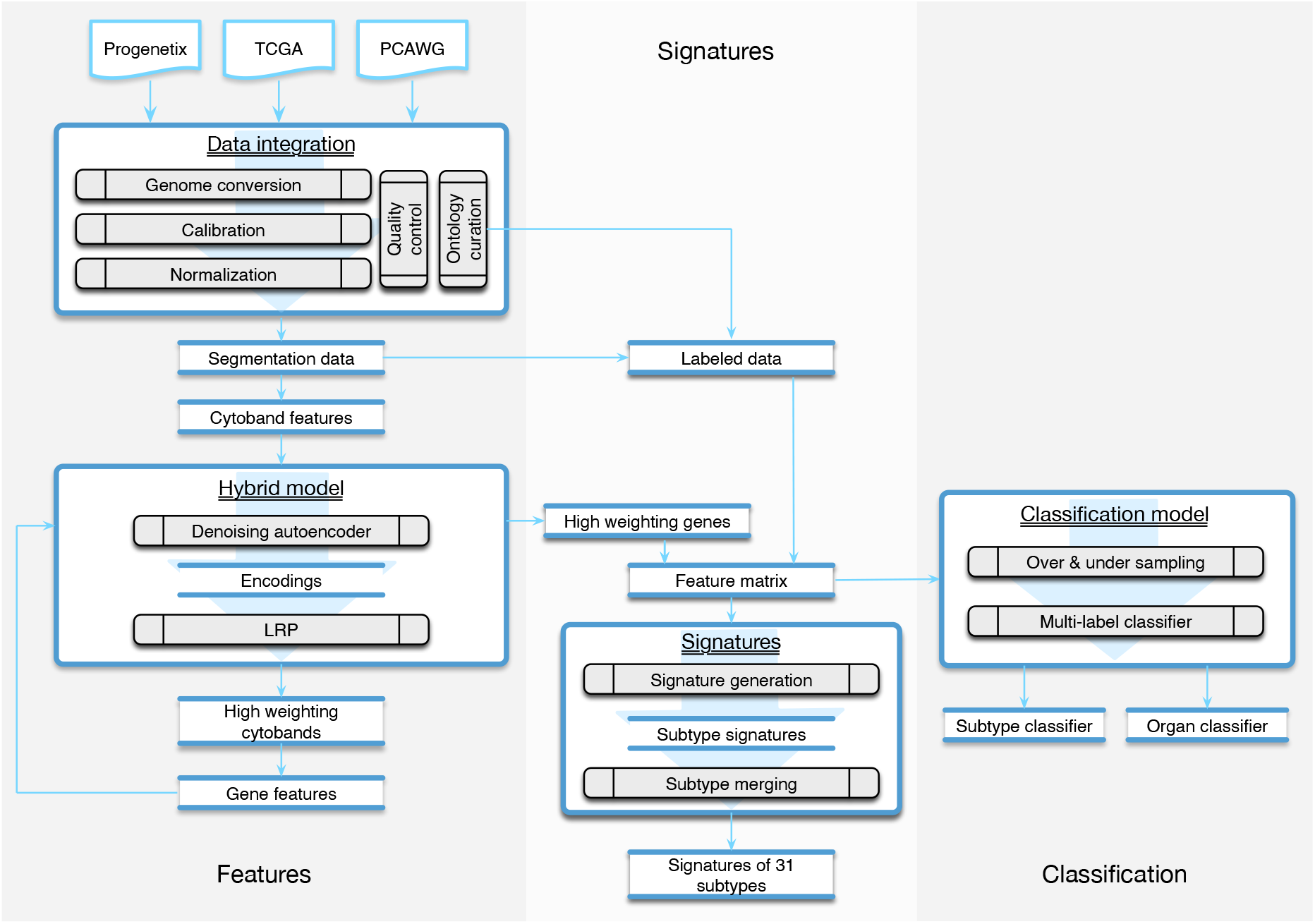
The workflow of the study was composed of three parts. The *Features* part consisted of methods of data integration and feature generation. The *Signature* part focused on creating CNA signatures for cancer subtypes and the categorization of subtypes. The *Classification* part recruited machine learning techniques to predict the organ and the subtype from a given copy number profile.

## Method

### Data integration

In this study, we integrated CNA profiles of tumor samples from three prominent resources. *Progenetix* [38] is a curated resource targeting copy number profiling data in human cancer. It features a large collection of data from different tissues and platforms. *The Cancer Genome Atlas (TCGA)* [39] represents a comprehensive cancer genomics initiative that generated standardized molecular profiling data for a wide range of cancer types. The *Pan-Cancer Analysis of Whole Genomes (PCAWG)* represents an initiative from the International Cancer Genome Consortium (ICGC) for the uniform analysis of a representative set of cancer samples using Whole Genome Sequencing (WGS) [40]. All three repositories provide publicly - though with the exception of Progenetix - partially access-controlled genomic data together with clinical information and diagnostic classifications. The initial collection consisted of 56,077 samples, representing an accumulation of CNA data from a wide range of tumor types and technological platforms.

When integrating data of high diversity for analyses, two major technical challenges can be encountered. First, genomics studies performed over the last decade will have applied different reference genome editions in their analysis pipeline, leading to shifted coordinates in results such as CNV segments. It is crucial to convert all data to the same coordinate system when analyzing data from multiple studies through a remapping procedure. However, unlike with SNP or other types of mutations involving short DNA sequences, applying standard remapping tools such as liftover [41] would result in a considerable information loss when converting copy number data, due to the occurrence of disruptive remapping in a relevant proportion of CNA segments. We previously addressed this problem with the development of a generic tool named *segmentLiftover*, to convert CNA data with high efficiency and minimized data loss [42]. Second, regardless of the underlying technology, genomic copy number data is derived from the relative assessment and integration of multiple signals, with the data generation process being prone to contamination from several sources. Estimated copy number values have no absolute or strictly linear correlation to their corresponding DNA levels, and the extent of deviation differs between sample profiles, which poses a great challenge for data integration and comparison in large scale genome analysis. To tackle this problem, we designed a method called *Mecan4CNA*(Minimum Error Calibration and Normalization for Copy Numbers Analysis) to perform a uniform normalization [37].

In the integration pipeline, samples were first converted to GRCh38(Hg38) using *segmentLiftover*. Then, the values in the sample were aligned to the corresponding true copy number levels of the main tumor clones using *Mecan4CNA*. For each step, a quality control protocol was set up to filter out samples carrying a low number of alternations, causing ambiguity in interpretation, or such having incomplete diagnostic information. The details of the integration are elaborated in Supplementary Implementations. In the end, a normalized dataset of high-quality samples was generated, which consisted of 44,988 tumor samples covering 271 ICDO-topography terms and 211 ICDO-morphology terms (Table 1).

**Table 1:**
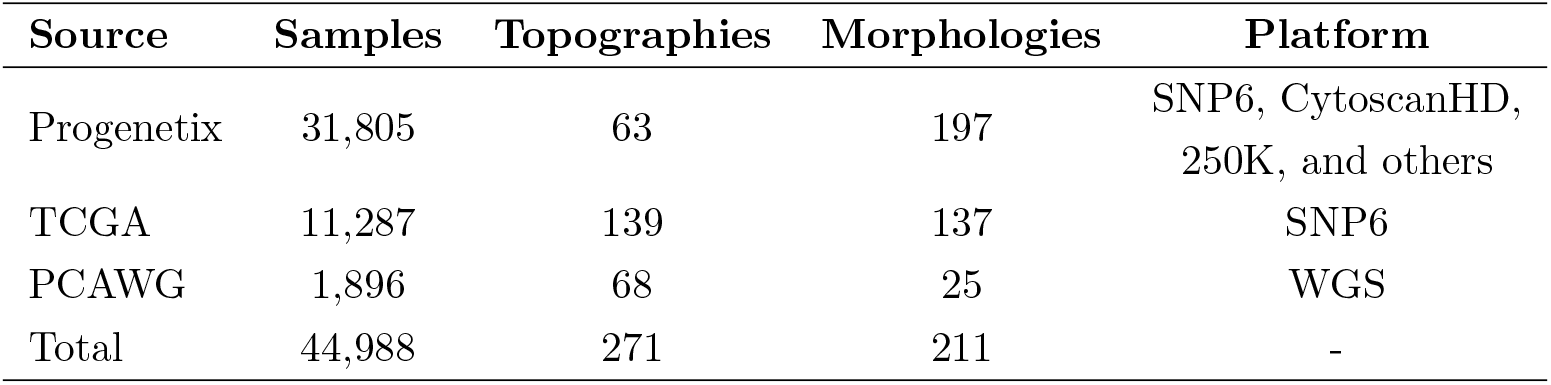
The data composition in the final dataset after quality filtering.

| Source | Samples | Topographies | Morphologies | Platform |
| --- | --- | --- | --- | --- |
| Progenetix | 31,805 | 63 | 197 | SNP6, CytoscanHD,<br>250K, and others |
| TCGA | 11,287 | 139 | 137 | SNP6 |
| PCAWG | 1,896 | 68 | 25 | WGS |
| Total | 44,988 | 271 | 211 | - |

### Feature representation

All copy number data can be represented in a platform-independent format of genomic segments, where each segment represents a continuous region on a chromosome. As the segments vary dramatically in platform-dependent resolution, reported size, and biologically supported thresholds [43, 44], a pragmatic approach for cross-study normalization lies in the creation of equally sized genomic bins and mapping of segment data into those. This method is elastic on the number of generated bins (features); therefore, it cooperates well with follow-up computational models that are sensitive to the feature numbers. However, the binning method disregards the uneven distribution of genes and does not capture any functional or structural information of the genome, thereby limiting biological correlations between features and the final model.

Another straightforward approach is to use the coverage of protein coding genes to represent each segment. This method provides inherent connections to genetic functions; however, direct mapping to protein coding genes overlooks the structural impact of CNA and also results in a large number of features, which are often difficult for machine learning methods to perform in optimal. In this study, we designed a two-phase approach to generate a refined gene panel as the feature representation, so that the noise and redundancy of the feature space could be reduced with consideration to functional significance. In the first phase, CNA segment data was mapped to cytobands, which represent “genomic phenotypes” correlated to gene density and chromosome structure and provided a good balance between feature number and genomic resolution. The cytobands features were then added to a hybrid model of *Autoencoder* and *Layer-wise Relevance Propagation (LRP)* to perform a feature extraction, which generated a panel of cytobands with high separation power. In the second phase, the selected cytoband features were denormalized to the protein coding genes from each band. Finally, another iteration of the feature selection process was performed to arrive at a selection of genes that contribute to the distinctiveness of individual sample profiles.

### Feature extraction

#### Autoencoder

An autoencoder is an unsupervised neural network that can derive abstracted representations from data. It usually consists of two parts, an encoder and a decoder, and both are also neural networks on their own. The input is first transformed to an encoding, then restored to its original by the decoder. The aim of an autoencoder is to reconstruct the input as exact as possible, and in the process, it learns a representation (encoding) of the input. It’s typically used for dimensionality reduction and to remove noise from signals [45, 46, 47].

An autoencoder possesses several features that are suitable for our research. Its ability to remove noise is well suited to limit the amount of “noise” from true passenger mutations, which may represent a considerable part of the copy number variations in tumors but do not have a functional impact. In the process of restoration, the encoding is able to capture the traits and distinguishing details, which is ideal for sample characterization in terms of their individual uniqueness. Finally, as a neural network, it can benefit from the extensive collection of heterogeneous data through its generalization capability. In this study, four different autoencoders (basic, denoising, sparse and contractive autoencoders) were compared on their abilities of input reconstruction. The denoising and contractive autoencoders showed the best performance in the restoration of CNV data, and the denoising autoencoder was chosen for its efficiency in implementation. The details of the comparison are elaborated in Supplementary Implementations.

#### Layer-wise Relevance Propagation

Layer-wise Relevance Propagation (LRP) is a technique to scale the importance and contribution of input features in a deep neural network. It utilizes the network weights and activation functions to propagate the output backward until the input. The basic propagation rule is illustrated as the following:

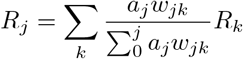

Here, *j* and *k* are two neurons from two adjacent layers. *a* is the activation of a neuron, and *w* is the weight between two neurons. The initial *R* is known from the output, and this formula is iterated to compute *R* for each neuron in the previous layer until reaching the input layer. LRP provides a solution to the “black-box” problem of deep neural networks, and is especially useful in tracing the network attentions and understanding the network behaviour. In our hybrid model, we exploited LRP’s ability to quantify the importance of input features in distinguishing different samples. When an autoencoder can reconstruct a large number of inputs with high accuracy, it provides an encoding that carries the information to recognize the differences between samples. If we apply LRP to this encoding, we would be able to measure the importance of each feature in the encoding.

#### The hybrid model

A primary goal of the research was to investigate the similarity and differences of CNA between cancers, through CNA feature dimension reduction towards an optimal representation of the uniqueness of each sample. In our hybrid model, we combined autoencoder’s ability to recognize differences and the LRP’s ability to scale feature relevance. First, the transformed feature matrix was used as the input data for the training of a denoising autoencoder. Then, the encoding generated by the autoencoder was used as the input of LRP to compute the weightings of each feature in the initial feature matrix. Next, the weightings of each feature were summed and normalized for all samples. Finally, by applying a threshold, we obtained a panel of high weighting features representing the uniqueness of each input sample.

In the two-phased feature extraction, the hybrid model first returned a panel of *cytobands* with high weighted CNA coverage; then, the hybrid model was utilized on the new feature matrix (CNA coverage on protein coding genes of the high-weighting cytobands), to calculate a panel of high-weighting *genes* as a representation of each sample’s feature space. Table 3 shows the resulting feature numbers for each step. The details of the procedures are provided in *Supplementary Implementations*.

### Signature generation

#### A dataset of major cancer types

To derive the genomic characteristics of different cancer types, it is important to utilize accurate, systematic mappings between samples and disease classifications. Following the standardized protocol set up for the *Progenetix* and *arrayMap* resources [38, 48], all samples were labeled using morphology and topography defined by International Classification of Diseases for Oncology (ICD-O 3) [49]. The complete data collection included 211 unique morphology (ICDOM) and 271 unique topography (ICDOT) terms. While the combination of ICDOM and ICDOT terms in principle can result in detailed disease classifications, the granularity of available sample descriptors varied dramatically between different studies. For example, while some samples were annotated as “epidermoid carcinoma, keratinizing” at “lower lobe, bronchus”, for other pulmonary carcinomas a more general “squamous cell carcinoma” of the “lung” was available in the input data. Also, the number of samples mapped to individual classification terms was extremely imbalanced. For example, while “infiltrating duct carcinoma of breast” was available with more than 5000 samples, “mucinous adenocarcinoma of lung” only was represented with 14 samples. For the sake of statistical validity and to minimize small-batch effects, we decided to focus on a data subset of 11 major organ systems with abundant samples and reliable copy number mutation content.

#### Signatures of cancer subtypes

After the two-phase feature extraction, we obtained a panel of genes, which drastically reduced the feature space to the signals of 927 genes. For any given sample, a small subset of the genes from this panel reflects its “signature” mutations and is able to represent the uniqueness of the sample against all other samples. When aggregating the samples of a disease type, we should be able to identify a subset of feature genes that are significantly altered in this disease (implementation details in Supplementary Implementations). Thus, we could use this subset of feature genes and their intensities as the CNA signature of the group. However, as mentioned in earlier sections, the samples were not labeled with the same granularity. For the clarity of analyses, samples of similar signatures were grouped together with a consistent label. First, samples were grouped into subtypes by the combination of their topography and morphology. Subtypes with less than 50 samples were removed. Then, the signature of each subtype was generated and the Pearson’s correlation coefficient of signatures was computed for subtypes of the same organ. Next, for each organ, subtypes of the same morphology level were merged into a more general morphology term if their correlation is greater than 0.9. Similarly, a subtype with a more detailed morphology is merged into another subtype of a more general morphology if their correlation is greater than 0.9. Finally, a dataset with refined disease labels was produced, which consisted of 22,671 samples covering 11 topography (organs) and 31 cancer subtypes. Table 2 shows the number of selected samples and subtypes in each organ. The complete table of subtypes and the merging are presented in Supplementary Subtypes.

**Table 2:** The number of samples and cancer subtypes in the studied organs.

| ICDO topography | Samples | subtypes |
| --- | --- | --- |
| Breast | 6,255 | 3 |
| Lung | 3,968 | 5 |
| Brain | 2,808 | 5 |
| Ovary | 2,128 | 3 |
| Colon | 1,832 | 4 |
| Kidney | 1,385 | 2 |
| Stomach | 1,303 | 5 |
| Skin | 1,238 | 1 |
| Prostate | 1,198 | 1 |
| Cerebellum | 1111 | 1 |
| Liver | 673 | 1 |

**Table 3:**
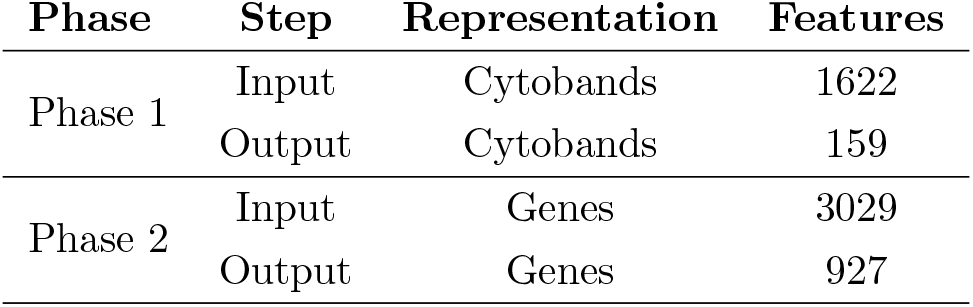
The number of features before and after each extraction phase. Cytoband features represent CNA occurrence on cytoband locations. Gene features are CNA coverage of protein coding genes, on the selected high weighted cytobands.

### Classifiers of subtypes and organ involvement

The compact feature panel and the curated labels provided us an opportunity to apply machine learning methods of classification performance. We constructed a multi-label classifier using the signature genes and their normalized signal intensities as features, and the subtype of each sample as labels. The primary challenge of building a multi-label classifier came from the extremely imbalanced number of samples in different classes, because the classifier would be significantly biased by the classes with a high number of samples. To create a robust classifier, we first applied undersampling on classes with a high number of samples and oversampling on classes with a low number of samples, so that the number of samples in each class became relatively balanced. Then, we recruited a random forest to learn and predict the subtypes. In addition to the prediction of subtypes, we were also able to evaluate the classifier’s performance on predicting samples’ organ of origin by mapping the subtype labels from predictions to the corresponding organ labels. Comparing with the approach that directly used the organ of each sample as labels in training, the subtype mapping method showed an improved performance. The implementation details of the model and comparisons are presented in Supplementary Implementations.

## Results

### CNA signatures

By using the data processing and modeling procedures described in the *Method* section, we generated a panel of feature genes from the collected CNA samples. Then, with the feature genes, we were able to create an abstract representation for each copy number profile, where only alternations that contributed to the distinctiveness of the sample were preserved. Figure 2 compares the original CNA patterns with the derived signature features where the frequent and extensive regional alterations in the original data have been replaced by a small number of feature genes, which visibly compare to subsets of characteristic changes in the original CNA data and represent the most discriminative alternations. With our methodology, ubiquitous alterations such as deletions on the short arm of chromosome 8 (Figure 2) are considered “non-typical” and therefore are not represented in the abstracted signatures.

**Figure 2:**
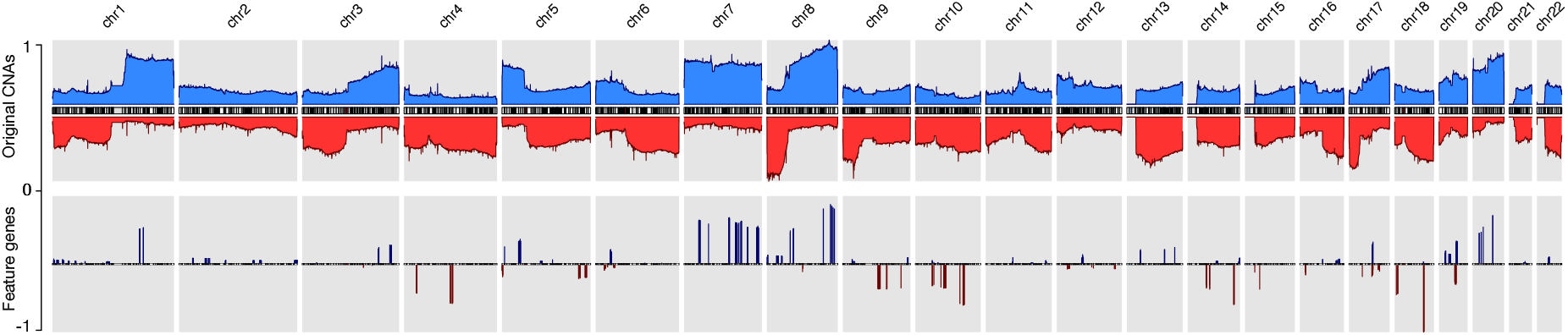
The copy number alternation landscape of all samples using the original CNVs and the feature genes. The feature genes are able to dramatically reduce the complexity of CNA signals while maintaining the mutational characteristics. The blue colors above the chromosome axis represent the average amplifications, and the red colors below the chromosome axis represent the average deletions. The amplitude of amplifications and deletions are normalized to [0,1] separately.

While the complete panel of feature genes provides the feature space for the whole of analyzed samples, in each individual sample only a small subset of these features reflects the sample’s own mutations. By aggregating samples from individual cancer subtypes, we were able to deduct the set of subtype related, significantly altered feature genes and could generate the CNA signatures of 31 cancer subtypes. Every signature consists of a subset of the general feature genes and their relative signal intensity comparing to other signatures. Figure 5 illustrates parts of the signature of *medulloblastoma*, *glioma* and *melanoma*. The complete list of signatures in all subtypes are included in *Supplementary Signatures*.

In the majority of copy number studies, analyses of tumor samples are focused on identifying the driver genes or the focal regions. However, in this study, the frequent drivers such as TP53 and PTEN, and the common aberrations such as CDKN2A/B and MYC are removed due to their prevalence. The genes in the signatures rather reflect the uniqueness of each sample or each cancer subtype. It is importance to note that the feature genes are not intended as the only nor the optimal representation. The fundamental objective of the study is to explore the potential driver mutations that are infrequent but relevant to specific cancer types. Although the signatures do not imply pathogenic causation, we can instead reveal their potential implications and correlations by investigating the signatures and the feature genes (Figure 6). In general, spatial and annotation analysis suggest that some feature genes reflect structural and functional alternations in samples; the signatures of different subtypes show high preference in several genomic regions that suffer frequent CNAs in many cancer types; and some subtypes, which have distinct disease codes, exhibit high similarity in their signatures.

### Focal and regional feature groups

The vast number of CNAs in cancer distribute in a wide spectrum of sizes, ranging from several kilobases to entire chromosomes. Recent research showed they are neither randomly nor uniformly distributed [3, 50]. Regarding their individual extent, CNAs can be categorized into two primary groups: they are either focal, affecting a region with a limited set of potential target genes, or very large and covering an extensive fraction of a chromosome. It has been found that - in general - these two groups of CNAs are caused by different biological mechanisms and can play different roles in tumorigenesis [51, 52, 53]. Focal CNAs usually arise from errors during DNA repairs and can suppress or promote the affected genes. On the other hand, chromosome level CNAs usually arise from errors during mitosis and can lead to dramatic structural aberrations, also involving large numbers of genes. Therefore, it seems important to explore the distinctions of focal and larger CNAs among the signatures’ features.

In studies of focal and large CNAs, no strict consensus on exact size the limits exists though general practice is to use an upper limit of 1-3Mb for “focal” events [54, 44] while others used a proportional cut-off such as 90% of a chromosome arm for “chromosomal” CNA [3, 50]. Because the signature feature genes are the simplification of the original CNA landscape, all chromosome arm level variations have been reduced to much smaller representations. Instead of studying arm-level CNAs, we merged adjacent feature genes within the distance of 5 MB to construct regional features, which indicates the high frequency and significant discriminative weightings of the covered region and the nearby regions. Feature genes, which are away from others for more than 5 MB on both ends, were considered focal features. After merging, the 927 signature genes were separated into 15 focal features and 43 regional feature groups. Figure 3(a) and (b) show the distributions of the different feature groups on the entire genome by their types and sizes. As expected, there are more regional feature groups than focal features in the merged feature space in both amplifications and deletions. This is because high-frequency focal CNAs were removed in the process of feature selection for their weak discriminating abilities. Among the regional groups, there are 50% more small regions (less than 5 MB) than large regions, especially in the deletions. It suggests that the small regions may represent the difference of the original arm level alternations between different cancer subtypes. Although at a lower frequency, the size of large regional groups can extend up to 200 MB, which strongly suggests that they represent distinctive arm-level events at these loci. Spatially, the large regions often occur exclusively and do not overlap with the same or the other type of CNVs. As illustrated in Figure 3(c), the large regional groups are mostly of low amplitude (the average of normalized CNV log-ratios across all samples); and no significant correlations with the amplitude are observed between different groups or types. The analysis of regional and focal features’ patterns across a large number of samples and cancer types confirmed and agreed with observations in several previous studies[2, 3, 6, 28]. The size distribution of merged feature groups also provided important implications to the underlying representation of different features.

**Figure 3:**
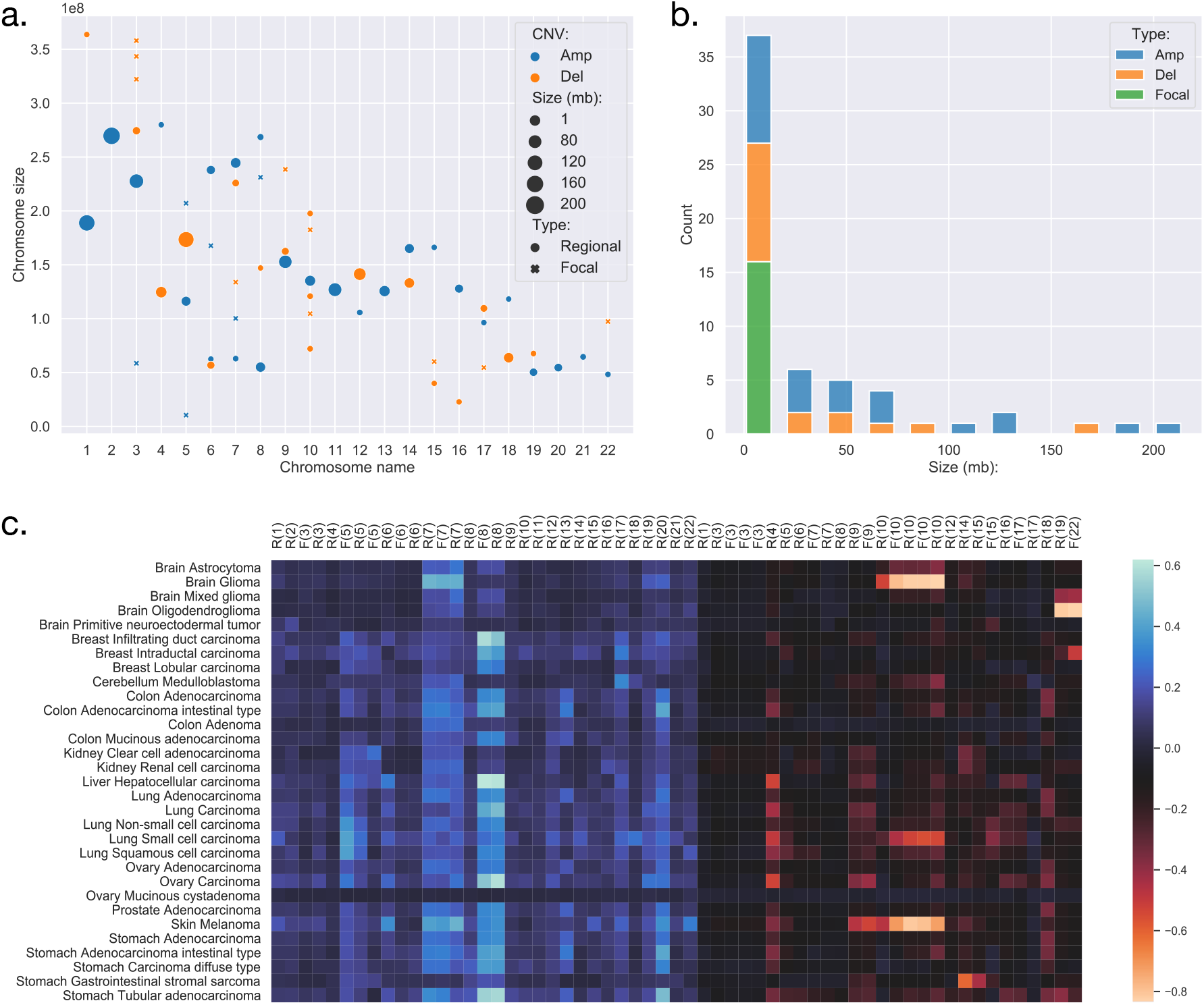
(a) The distribution of focal and regional feature groups on the genome. The groups were generated from the 927 feature genes, where genes within 5 million-bases were merged into a group. The position of the mark was determined using the centre of the group or gene. The size of the mark was determined by the length of the group or gene, in the unit of million-bases and a minimum size of 1 MB. (b) The length distribution of feature groups. (c) The amplitude of feature groups in 31 studied cancer subtypes. The values were computed using the average of the normalized CNV log ratios of all samples in each subtype. On the x-axis, F denotes focal, R denotes regional, the number denotes chromosome name.

### Functional implications of the feature genes

A cardinal concept in this study was to describe cancer subtypes using CNAs based on their uniqueness and discriminating ability, instead of their frequencies. The selected feature genes usually exhibited different patterns in either their presents or variation amplitude between different subtypes. As a result, the final features mainly included non-focal and less prevalent genes that do not play known central roles in tumorigenesis. However, it is still interesting to investigate the feature genes by their annotations and discuss their potential functional meanings.

The HGNC (HUGO Gene Nomenclature Committee) gene families are defined based on evolutional and functional homologies [55]. Table 4 lists the significantly over and under expressed HGNC families, where the family has less than 0.05 p-value in Binomial test and includes at least 5 feature genes: (1) The immunoglobulin heavy chains related genes showed the highest significance. Although the mutations of these genes are known to play causal roles in B-cell related malignancies [56, 57], surprisingly, accumulating evidence has also shown their frequent overexpression in epithelial cancers [58, 59]. Though their functions and mechanisms in oncogenesis are still not well understood, several studies have demonstrated their tumor-promoting impacts and observed their prognostic implications in breast cancer and ovarian cancer [60, 61, 62]. (2) Olfactory receptors are one of the largest gene families in the human genome. Similar to immunoglobulin genes, their overexpression has also been observed in various cancers and is often neglected in genomic cancer studies because of their specific role [63]. However, recent studies have begun to reveal their functional roles and prognostic potentials in prostate cancer and breast cancer [64, 65, 66]. (3) The tripartite-motif (TRIM) family proteins are involved in a variety of biological processes including transcriptional regulation, cell growth and apoptosis, and their dysregulation has been extensively linked to cancer risk and prognosis [67, 68, 69]. (4) The HOXL subclass homeoboxes are overexpressed in amplifications and underexpressed in deletions. The family consists mainly of HOX genes, which are master regulatory transcription factors in embryogenesis, cellular development, and issue homeostasis [70]. Recent researches have shown that depending on the type of the tumor, the expression of HOX genes may be increased or decreased and play roles in oncogenesis or tumor suppression [71, 72, 73]. (5) Keratin-associated proteins have no confirmed role in cancer. (6) The combined actions of glycosidases and glycosyltransferases constitute the primary catalytic machinery for the synthesis and breakage of glycosidic bonds. The aberrant glycosylation patterns are often considered a hallmark of cancer, which promotes tumor proliferation, invasion, and metastasis [74, 75, 76, 77]. (7) Intermediate filaments proteins are often used as diagnostic markers in cancer, because changes in their expression patterns are often associated with tumor progression and cancerous cells usually partially retain their original structural signatures [78]. Several recent studies have reported their active roles in tumor invasion and metastasis [79, 80]. (8) Zinc fingers C2H2-type family is the largest group of transcription factors in humans. A number of studies have revealed that aberrant expression of C2H2 proteins promotes tumorigenesis with various roles in several cancer types [81, 82].

**Table 4:** The significant HGNC gene families of the 927 feature genes, where the family is covered by at least 5 genes and with the Binomial test less than 0.05.

| **HGNC family** | **Type** | **P-value** | **Potential impact** |
| --- | --- | --- | --- |
| Immunoglobulin heavy locus at 14q32.33 | AMP | 3.18E-08 | Increased IGH expression, prognostic correlations |
| Olfactory receptors, family 52 | AMP | 6.83E-06 | Increased OR expression, prognostic correlations |
| Tripartite motif containing | AMP | 3.71E-04 | Transcription, cell growth, apoptosis |
| Olfactory receptors, family 51 | AMP | 7.10E-04 | Increased OR expression, prognostic correlations |
| HOXL subclass homeoboxes | AMP | 1.83E-03 | Oncogenesis, tumor suppression |
| Keratin associated proteins | AMP | 4.11E-03 | N/A |
| Glycoside hydrolases | AMP | 9.84E-03 | Invasion, metastasis |
| Glycosyltransferases | AMP | 3.26E-02 | Invasion, metastasis |
| Intermediate filaments | AMP | 3.44E-02 | Invasion, metastasis |
| Zinc fingers C2H2-type | AMP | 3.50E-02 | Transcription, multiple roles |
| ANTP class homeoboxes | DEL | 4.40E-05 | Oncogenesis, tumor suppression |

The functional background of the significantly expressed HCNC families reveals several exciting aspects of the feature genes. First, the features consist of biomarker genes, which are frequently and abnormally expressed in many cancer types. Although their exact mechanisms in tumorigenesis are not fully understood, many studies have confirmed their functional roles and prognostic correlations in cancer. Second, the features describe functional aberrations by including several families that are related to transcriptional regulations. Previous studies have shown the high correlation of CNAs and differential gene expression in cancers [83, 84], and partially elucidated their involvement in tumor development and progression. Finally, the features include hallmark signatures in invasion and metastasis, which are crucial indications of late-stage cancer progression. They are more likely to represent the consequence of other mutational activities rather than the direct impact of CNAs.

In summary, the analysis of feature genes agreed with a variety of previous research in their cancer-promoting or related roles and suggested the potential rationales behind the features. It also confirmed that malignant cell transformation is often an orchestrated process of many interplaying parts. While important driver mutations can signify the milestones in tumor development, the minor mutations during the development can also provide valuable characterization to specifics between different cancer types.

### Discriminative genomic loci

Since feature genes represented the uniqueness of individual samples, it is expected that they would distribute exclusively among different subtypes in general. Interestingly, while most of the features are sparsely distributed, there are also a few “hot zones”, which are frequent CNV regions in general, that are commonly included in many signatures with high significance. Figures in the *Supplementary Distribution Density* illustrate the distribution and frequency of duplication and deletion feature genes separately. Figures on the first row are plotted using the full panel of feature genes, and figures on the second row are plotted using feature genes from all signatures (i.e. significantly high frequency). The most prominent feature genes among subtypes are those representing duplications on chromosome 7 and 8. Specifically, the feature genes are distributed in groups in several regions: *chr7:28,953,358 - 34,878,332* and *chr7:49,773,638 - 50,405,101* on the p arm of chromosome 7; *chr7:91,692,008 - 93,361,123*, *chr7:105,014,190 - 117,715,971*, *chr7:129,611,720 - 130,734,176* and *chr7:148,590,766 - 152,855,378* on the q arm of chromosome 7; *chr8:47,260,878 - 50,796,692*, *chr8:51,319,577 - 54,871,720* and *chr8:112,222,928 - 138,497,261* on the q arm of chromosome 8. As shown in Figure 2, chromosome 7 and 8 contain the most frequently duplicated regions among all studied cancer types. Despite their common existence, analysis of individual signatures and samples reveals three patterns that may explain their prominence in features: first, on the two chromosomes, samples often have large duplications. The aggregated signals in individual subtypes usually show high alternation frequency of the whole region and pose a plausible hint of frequent chromosome or arm level events. Second, the spans of duplications differ among subtypes. For example, the signature of melanoma consists of feature genes covering the entire chromosome 7 and 8; the signature of lung adenocarcinoma consists of feature genes only on the p arm of chromosome 7, and the signature of ovary carcinoma consists of feature genes only on the q arm of chromosome 8. Third, the alternation amplitude of the feature genes also shows distinctive patterns among subtypes. The subtle difference in the scale and amplitude of CNV on chromosome 7 and 8 suggest that aberrations of these hot regions may play differentiating roles in different cancers.

Similar to duplications, the most common deletion features distribute on three regions on chromosome 18: *chr18:2,916,994 - 7,117,797* on the p arm; *chr18:58,481,247 - 60,372,775* and *chr18:69,400,888 - 70,330,199* on the q arm. Different from duplications on chromosome 7 and 8, the deletions on chromosome 18 show a similar pattern of high amplitude deletions on the entire chromosome 18, where feature genes on the p arm suggest a one-copy deletion on average, and feature genes on the q arm indicate frequent homozygous deletions. Interestingly, the deletion features on chromosome 18 are usually mutually exclusive to the deletion features on chromosome 10, which are under high pressure of chromosome level deletions in several subtypes. Previous studies have observed strong correlations of deletions on chromosome 18 and chromosome 10 to the progression of several cancer types. Other analyses have investigated their functional impacts and suggested the presence of several tumor suppressor genes [83, 85, 86, 87]. Another hot zone of deletion features locates on the q arm of chromosome 22 (*chr22:48,489,460 - 48,850,912*). High-level deletions are frequently observed and selected as features in many subtypes. Different studies have shown similar observations of frequent deletions on chromosome 22, however, the mechanism of the deletion is still obscure and the functional impact remains putative [88, 89].

Besides high prevalence, the feature “hot zones” also exhibit a higher copy number alternation level than average features. As shown in Figure 4, the average copy number alternation levels are far greater in these regions than in aggregated features: 56% higher in amplifications, and 70% higher in deletions. Furthermore, several hot zones and the associated cancer types, including the frequent amplifications near the centromere of chromosome 7, on the q arm of chromosome 8, and deletions closed to the telomere of chromosome 18, are in coherence with the general CNA patterns identified in previous pan-cancer studies [3, 6, 90], which suggests these regions are related to high CNA frequency. More specifically, some hot zones show correlations with nearby driver genes, which are frequently observed with high CNAs in cancers having high alternations in these hot zones. First, the high alternations level on chromosome 7 hot zones are in accordance with the overexpression of nearby EGFR, MET, and BRAF genes in glioma, lung cancers and melanoma. EGFR is a well-studied oncogene in lung cancers and gliomas, MET is also a frequent driver in lung cancers, and BRAF is a signature driver gene in melanoma. A variety of studies have reported the CNA amplifications at or near the hot zones, and investigated their driver roles in these cancers [91, 92, 93, 94, 90]. Next, the frequent but relatively low-amplitude hot zone on the q arm of chromosome 8 is correlated with MYC, which is frequently amplified in numerous cancers. The extensively amplified hot zones on the p of chromosome 8 may be associated with FGFR1 and IKBKB, which are often deregulated by amplification in lung, colon, and bladder cancers. Studies of copy number patterns in these cancers have shown the significant correlations of increased gene expressions and amplification at the hot zone regions [95, 96, 97]. Finally, the supporting evidence on deletion hot zones are mostly prognostic instead of functional confirmed, except on chromosome 10, where two frequent and high-alternation hot zones are closed to PTEN. Although it is one of the most common inactivated tumor suppressor genes in cancers, the signal amplitude of these zones is in high consistency with cancer types (gliomas, lung cancers, melanoma, and breast), where the loss of PTEN is especially frequent and crucial [98, 99].

**Figure 4:**
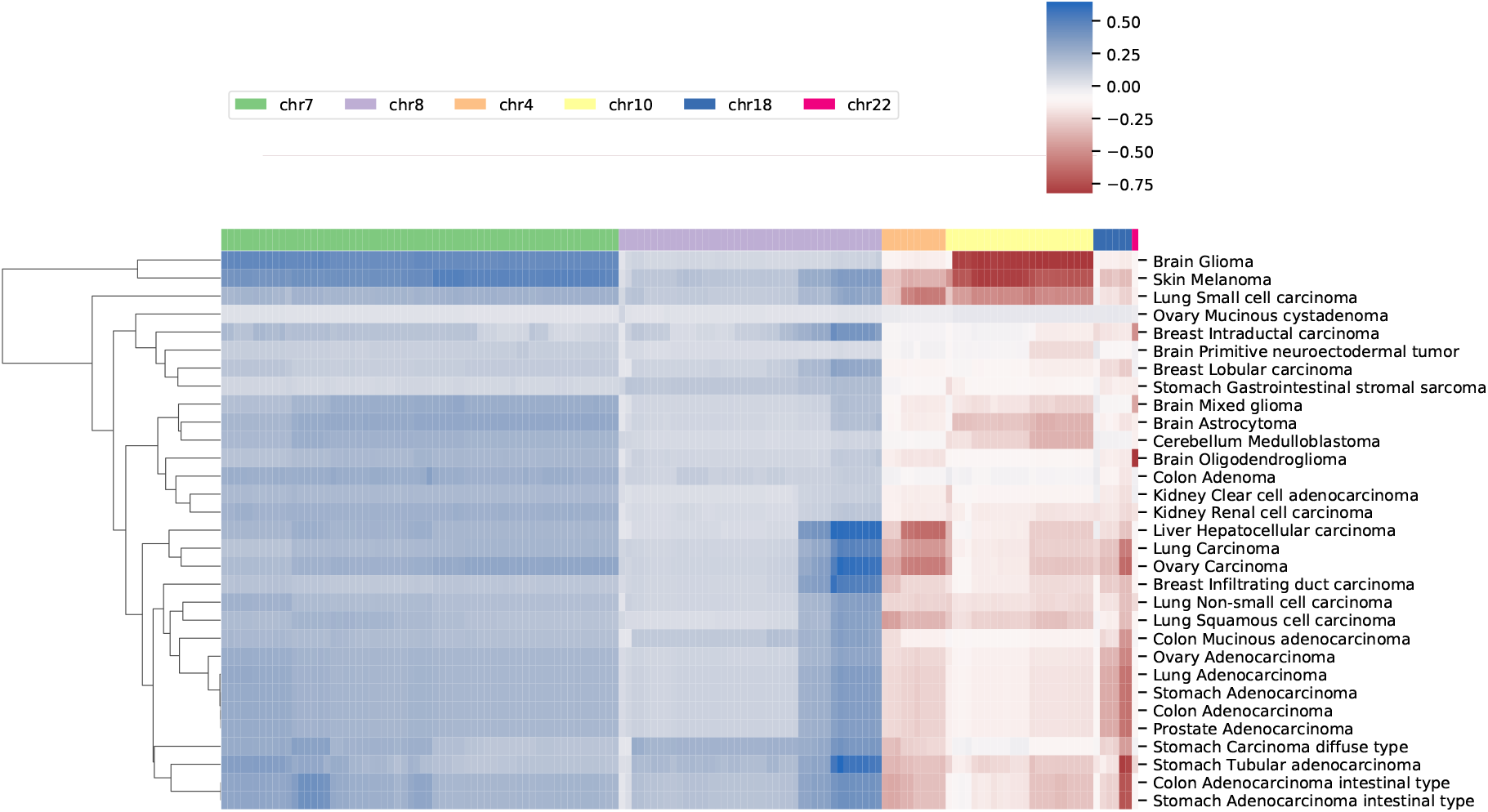
The average copy number alternation level of the hot-zone features among different cancer subtypes. The values were calculated using the mean of the normalized CNV log ratios of all samples in each subtype.

### Similarities of neural crest originated subtypes

The signature based clustering of cancer subtypes (Figure 6) shows that diseases in the same cluster usually also share close ontologies. However, there is an interesting outlier, which consists of medulloblastoma, melanoma and glioma (astrocytoma is a subtype of glioma). The three clinically and topographically distant cancer types have signatures of high similarity in both the selection of features and their alternation frequencies. Figure 5 shows the comparison of the original CNV data, the features and the known drivers of the three cancers on chromosomes harboring similar signatures. Their signatures exhibit high similarities in the duplication of chromosome 7 and the deletion of chromosome 10. They also share pairwise similarities on the duplication of chromosome 1 and 20, and the deletion of 9 and 14. For the three cancers, the most frequently amplified chromosome 7 harbors several of the key oncogenes. For example, EGFR, CDK6 and MET in glioma; KMT2C and PMS2 in medulloblastoma; BRAF, RAC1 and TRRAP in melanoma. The most frequently deleted chromosome 9 and 10 harbor several important suppressor genes. For example, CDKN2A and PTEN in glioma; XPA, PPP6c and CDKNA in melanoma; PTCH1 and SUFU in medulloblastoma. Noticeably, the CDKN2A/B deletion is the most frequent copy number aberrations across all cancer types. Although the distribution of driver genes shows a close correlation with the frequency and amplitude of CNV, most drivers do not demonstrate signal peaks nor overlap with the feature genes. Frequent duplications on chromosome 1 and deletions on chromosome 14 do not show direct correlations to common driver genes; however, a number of studies have shown the connection of CNAs on these regions to the progress and prognosis in these cancer types [100, 101, 102, 103, 104, 105].

**Figure 5:**
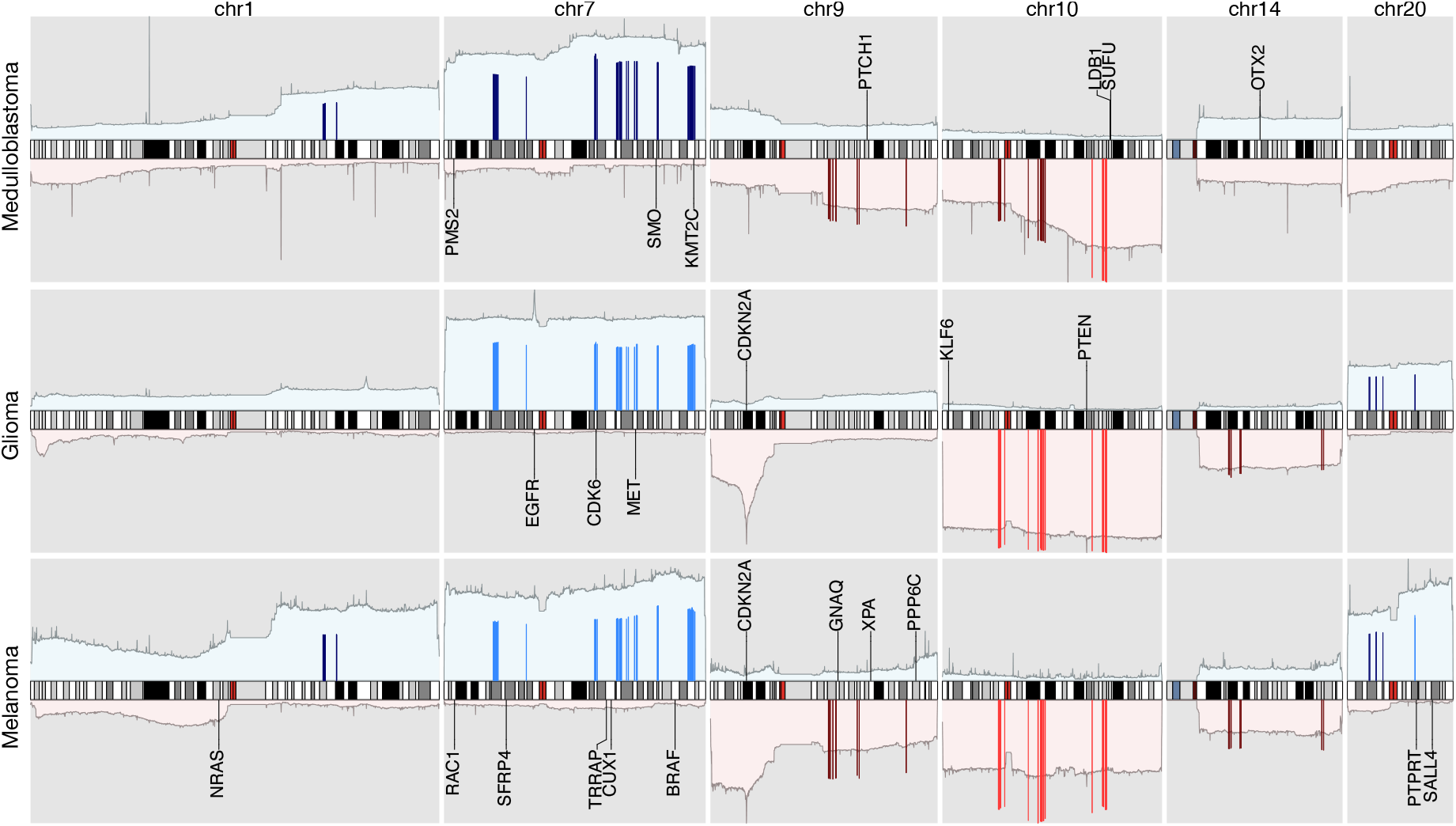
The integrated view of the original data and the selected features, in the neural crest originating entities medulloblastoma, glioma, and melanoma. The shaded background area color illustrates the original data. Color bars illustrate the feature genes, where brighter colors indicate stronger signal intensity. The blue colors above the chromosome axis represent the average amplifications, and the red colors below the chromosome axis represent the average deletions. The amplitude of amplifications and deletions are normalized to [0,1] separately. The adjacent known driver genes are also included for each tumor type.

**Figure 6:**
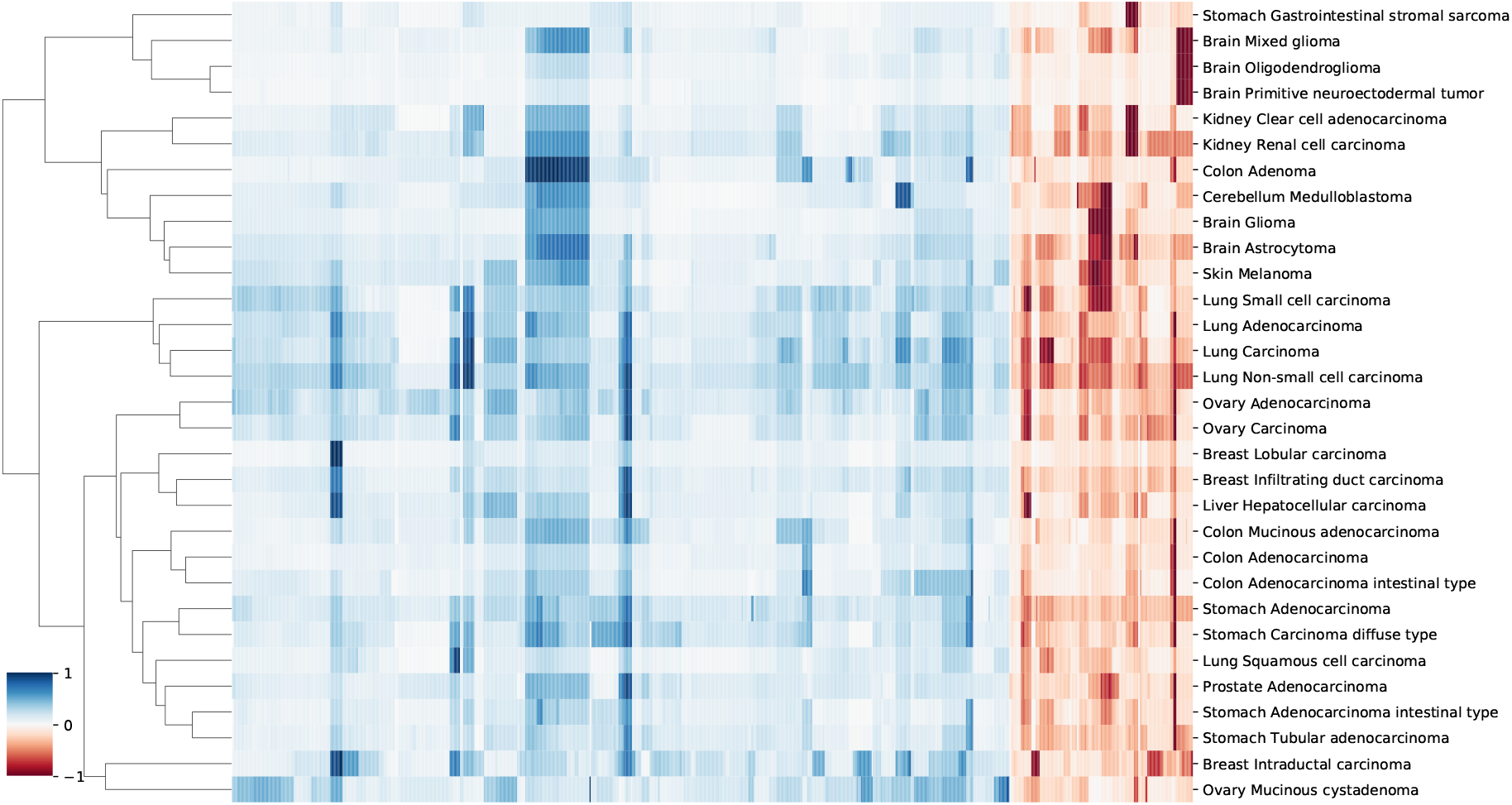
A clustering heatmap of features in 31 signatures. Columns are normalized average CNV intensities of feature genes, where the blue colours are duplication features and red colours are deletion features. Duplication and deletion frequencies are normalized separately.

In the 1990s, epidemic studies [106, 107, 108] first revealed the connection between melanoma and tumors of the nervous system: not only a familial association, which was confirmed by germline mutations; but also a significantly increased risk of one disease in people having a history of the other one. Although there was evidence showing their potential common pathophysiologic pathways and responsiveness to the same drugs, the genetic connection of the two disease groups was still largely unclear [109]. In spite of their ontological difference, medulloblastoma, melanoma and glioma are all derived from the lineages of neural crest cells. Recent studies of neural crest cells and the cancers from their lineage cells suggest that malignant cells mimic many of the behavioral, molecular, and morphologic aspects of neural crest development [110, 111, 112]. Aberrations in tumor cells may lead to the reactivation of their embryonic developmental programs and promote tumorigenesis and metastasis. For example, the WNT family members, which play import roles during the epithelial-to-mesenchymal transition of neural crest cell, are reactivated during invasive transformation in melanoma [113, 114]. In glioblastoma, experimental data suggests that the dysregulation of the WNT signaling pathway supports the onset of cancer stem cells, which assure the enlargement of the tumoral mass and eventually the spread of metastases [115]. In medulloblastoma, the WNT subgroup signifies one of the four molecular subtypes of the disease [116]. In regions represented by the similar signatures, several WNT genes exhibit abnormal amplification frequencies, which are a potential reflection of the overexpression of WNT signaling. Specifically, WNT2B and WNT4 are covered by amplifications of moderate frequencies, and WNT2, WNT3A, WNT9A and WNT16 are encompassed by amplifications of high frequencies. WNT2 and WNT16 are signature genes in all three subtypes implicating their prevalence among individual samples.

Another interesting result from the similar signatures is the connection between melanoma and medulloblastoma, which has dramatic differences in many biological aspects. For example, medulloblastoma is considered primarily originated from embryonal cells in early development. It mostly occurs in children and usually has good prognosis outcomes. On the other hand, melanoma is primarily caused by ultraviolet light exposure. Its risk has a positive correlation with age and the prognosis is usually very poor once passing the early stage of the disease. However, besides their differences, all three cancer types are notorious for their fast progression, high invasiveness and wide metastasis. It’s possible that their common CNA signatures are the imprint of their acquired cancer hallmarks instead of their mutational characteristics. The combined evidence suggests that different types of tumor may achieve their hallmark abilities through different evolution paths [117]. The acquired hallmarks exhibit a number of common genotypes such as copy number aberrations, which are potentially the downstream result of primary mutation events that contribute to the functional sustainability of tumor cells.

### Classification of subtypes and organs

In the previous sections, we explored the implications of the feature genes and cancer signatures. It is also interesting to assess the robustness and usefulness of the feature genes in sample classifications, especially when most of the primary driver genes are excluded from the features. The identification of the origin of a tumor sample is a challenging task that attracted significant attention from both academia and industry. A great number of computational methods have been proposed in recent years to facilitate the early diagnosis of cancers from liquid biopsies [29, 34, 118, 119, 120]. Among the various explored genomic aberrations, several studies have illustrated the predictive effectiveness of copy number aberrations [34, 35, 36]. In the meantime, machine learning researchers have also proposed a number of classification methods to predict cancer types using copy number data [121, 122, 123]. However, these methods usually utilized a uniform dataset such as TCGA to investigate the feasibility of the method and the performance of the model. They often lack the generalization experiments to apply the method on data that is not from the training data pipeline. Also, the goal of machine learning researches usually focused on constructing a classifier of high accuracy, whereas in this study, we focused on exploring the unique and typical mutations in different cancer subtypes. Therefore, we would like to use the classification results of copy number signatures to demonstrate their potential in the predictions of cancer subtypes and organs of origin from liquid biopsies, and performed no direct comparisons with other predictive studies.

In the prediction of subtypes, the model showed a wide range of performance spectrum on different subtypes. Figure 7(a) shows the confusion matrix of individual subtypes, where the x-axis represents the true labels and the y-axis represents the predicted labels. Specifically, the classifier achieved good performance in predicting: breast intraductal carcinoma, colon adenocarcinoma, brain glioma, cerebellum medulloblastoma, cerebellum medulloblastoma, and kidney clear cell adenocarcinoma, where the F1-score of individual subtypes ranged from 0.72 to 0.67. On the other hand, the prediction performance was deficient in several subtypes, such as brain astrocytoma, stomach carcinoma diffuse type, and colon mucinous adenocarcinoma. The complete metrics of each subtype are included in Supplementary Performance. Investigation on false negatives and false positives showed that most of the false predictions fell into the subtypes that were from the same organ and had significantly more samples. In general, the performance of individual subtypes was positively correlated with the number of samples (Figure 7(c)). The difficulty in predicting some subtypes suggests that some clinico-pathologically closed cancer subtypes share similar copy number mutation patterns. While the copy number signatures showed promising predictive potential for some cancer subtypes, data from complementary genomic aberrations are needed for a comprehensive model to classify similar subtypes.

**Figure 7:**
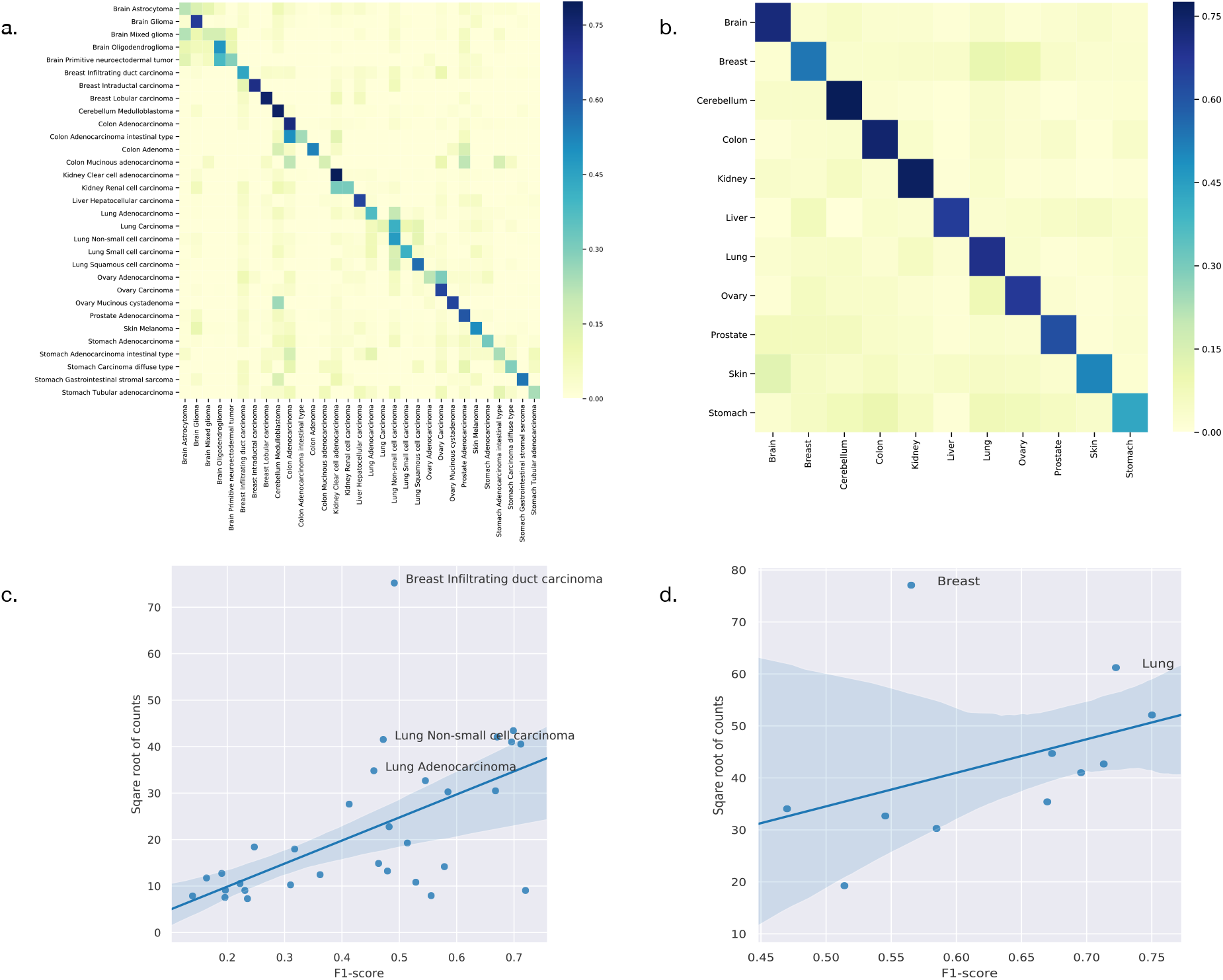
(a) The classification performance of individual subtypes. (b) The classification performance of individual organs. For (a) and (b), the x-axis are true labels, and the y-axis are predicted labels. (c) The correlation between the performance and the number of samples in the classification of subtypes. (d) The correlation between the performance and the number of samples in the classification of organs.

In the prediction of organs, the model showed a more consistent performance than in subtypes. Figure 7(b) shows the confusion matrix of individual organs. Specifically, the classifier achieved good results in predicting: brain, lung, colon, cerebellum, ovary, and kidney, where the F1-sore of individual organs ranged from 0.75 to 0.67. Although some organs showed relatively low prediction accuracy, they all had dramatic improvements compared with the prediction results of the divided subtypes. The complete metrics of each subtype are included in Supplementary Performance. In general, the performance was also positively correlated with the total number of samples in each organ (Figure 7(d)). In contrast, the prediction results of organs showed a significant overall improvement over the subtypes. For example, none of the lung subtypes had a good performance in the subtype classification. However, when combined, the lung achieved the second-best performance (0.72 F1-score) in organ prediction. A similar improvement was also observed for the stomach while the prediction performance of breast had a drastic decline compared with divided subtypes; the latter is further discussed in the subsequent section.

In summary, our classification experiments indicated that the copy number signatures have good potential in predicting several cancer subtypes and organs of origin. However, not all investigated subtypes and organs achieved satisfying performance with solely CNA features, which confirmed the need for ensemble strategies to identify unknown tumor cells’ type and origin. Moreover, the improved performance in organ classifications suggested the correlations of CNA and organ’s functional influence in tumorigenesis. In future research, it would be interesting to further investigate the difference between organ-specific CNAs and subtype-specific CNAs.

### Heterogeneity within subtypes

The results of the classifier showed significant differences in performance among different subtypes and organs. While some of the under-performance may be attributed to the insufficient number of samples, the false predictions of several classes suggested the heterogeneity within these subtypes. As shown in Figure 7(a), for breast infiltrating duct carcinoma and cerebellum medulloblastoma, the false predictions were widely distributed over several unrelated classes. Figure 7(c) and (d) highlighted breast infiltrating duct carcinoma, lung non-small cell carcinoma, and lung adenocarcinoma as outliers of the correlation, where the high number of samples did not improve but deteriorate their performance.

Large-scale cancer genome studies have shown that tumors from the cerebellum, lung, and breast usually carry high burdens of copy number mutations [6, 3]. With the extensive number of CNA in these subtypes, it is expected to be more challenging to identify key signals. Genomic studies in the past two decades have revealed mutation characteristics in these diseases, and established classifications of their molecular subtypes. For example, medulloblastomas are now commonly categorized into four subgroups: *WNT*, *SHH*, *G3* and *G4*. Similarly, breast carcinomas can be categorized into four subgroups: *Luminal A*, *Luminal B*, *Her2-enriched* and *basal-like*. Studies of their mutational patterns showed distinct markers and different mutation landscapes of each subtype, and multi-omics studies further revealed the intrinsic complexity within subtypes and suggested classifications of more subgroups [124, 125, 126]. However, the majority of the samples in our dataset were labeled by their morphology, with the molecular subtype classification missing in the sample annotations. As a result, large diagnostic groups such as “breast infiltrating duct carcinoma” could not be further separated *a priory*, resulting in a common label for breast carcinomas with the diagnostic subtype consisting of 5,657 samples, over three times more than the second-largest subtype. While we addressed these skewed overall subtype sizes through the application of undersampling techniques, the implicit mixture of different molecular subgroups provided an inevitable impact on the association of diagnostic labels and CNA based classifiers.

## Discussion

For this study, we assembled a large collection of cancer CNA profiles with the aim to identify genomic aberration signatures with specificity for individual cancer types. In order to integrate technically diverse data, we developed innovative tools for genome data conversion and signal normalization. Targeting the identification of unique components in diagnosis mapped CNA profiles, we were able to derive the CNA signatures of 31 cancer subtypes, where each signature was characterized by a minimal representation of genes with high discrimination capacity. The signatures were further evaluated to derive classifiers identifying tumor types and organs of origin. The comparative analyses of signature genes and their represented regions showed that duplications on chromosome 7 and 8, and deletions on chromosome 22 were the most common aberrations among studied cancer types. However, these regions also harbored features of high differentiating power, which may indicate the functional significance of cancer related genes with specificity for cancer-type related pathway involvement.

While our feature genes exhibited certain correlations with known driver genes, this itself does not provide direct evidence of their functional significance. Feature genes could also be the result of the downstream effects of mutational activities or be co-opted as a representation of an aberrant region where the pathogenetic activity is provided by different genomic elements [127]. In the analysis, we were prudent to limit propositions of the impact of individual feature genes.

Our analyses showed that three clinico-pathologically distant cancer types - medulloblastoma, melanoma, and glioma - shared CNA signatures of high similarity. Developmentally, the three tumor types can be traced back to common lineages of neural crest cells. Research of neural crest cells and epidemiologic studies of glioma and melanoma have shown sporadic evidence of their connections in cancer development, but the genetic foundation of their association with respect to oncogenetic processes is still elusive. Here, comparative analysis of shared mutations together with improvements of developmental processes in the corresponding normal tissues may provide insights into shared pathologies and potential therapeutic targets.

The studies on genomic classifiers demonstrated the capacity and potential of CNA signatures in tumor identification. However, while the multi-label classification approach showed promising performance in many cancers, in a few subtypes the performance was deteriorated by the intrinsic heterogeneity of their CNA profiles. Here we expect that future follow-up studies with extended input data and employing different ontology-based grouping strategies may lead to improved performance and the potential emergence of better group aggregations.

An essential challenge in the study was the integration of data from varying genomic profiling platforms. During normalization and quality control, we removed samples that were ambiguous in interpretation or had an abnormal distribution of signals. However, as a trade-off, many genomically heterogeneous samples (e.g. aneuploid or containing CTLP) were removed in the process. For future studies, complementary strategies should be developed to include these samples without detrimental effects on data normalization and quality. With respect to sample annotation for histologic and diagnostic classifications, extensive efforts went into the curation of the data with the aim to use a unified set of classifications. Here, future analyses may make use of emerging cancer specific ontologies for a better integration of varying annotation granularities using hierarchical terms and concepts. Currently, our research group is finishing a parallel study on ontology mappings of cancer samples, and we are expecting a considerable improvement in the disease annotation of samples in future studies.

In summary, this study presents a systematic pipeline for integrative and comparative analyses of a large amount of copy number data. The resulting CNA signatures offer new perspectives on the understanding of common foundations in cancers and show promising potential in applications of tumor classification.

## Supporting information

Supplementary implementations

Supplementary Density Plot

Supplementary Normalization Examples

Supplementary Performance

Supplementary Signatures

Supplementary Subtypes

## Author Contributions

BG developed the method and implemented the analysis. MB conceived the original concept and supervised the project. Both authors contributed to the writing and revisions of the manuscript.

## Acknowledgments

We thank Paula Carrio Cordo and Qingyao Huang for support with the data ontologies, and all current and previous members of the Baudis group for contributions to the Progenetix resource.

## Supplemental Data

- Implementation details: Supplementary Implementations.pdf
- Performance details of individual classifications: Supplementary Performance.pdf
- Analyses results of subtypes: Supplementary Signatures.zip
- Labeling and merging of subtypes: Supplementary Subtypes.pdf
- Distribution and amplitudes of feature genes in different subtypes: Supplementary Distribution Density.pdf
- Illustration of cross-sample CNV normalization: Supplementary Normalization Examples.pdf

## Data Availability Statement

- Segmentation data and metadata of open-access samples used in the study are available at https://progenetix.org/gao-2021-signatures/search/
- Data from TCGA is available at https://portal.gdc.cancer.gov
- Data from PCAWG is available at https://dcc.icgc.org
- Development repository is available at https://github.com/baudisgroup/cancer-signatures

## References

[1] Zhang F, Gu W, Hurles ME, Lupski JR. Copy number variation in human health, disease, and evolution. Annual Review of Genomics and Human Genetics. 2009;10:451–481.

[2] Baudis M. Genomic imbalances in 5918 malignant epithelial tumors: an explorative meta-analysis of chromosomal CGH data. BMC Cancer. 2007;7(1):226.

[3] Beroukhim R, Mermel CH, Porter D, Wei G, Raychaudhuri S, Donovan J, et al. The landscape of somatic copy-number alteration across human cancers. Nature. 2010;463(7283):899–905.

[4] Cox C, Bignell G, Greenman C, Stabenau A, Warren W, Stephens P, et al. A survey of homozygous deletions in human cancer genomes. Proc Natl Acad Sci U S A. 2005;102(12):4542–4547.

[5] Upender M, Habermann J, McShane L, Korn E, Barrett J, Difilippantonio M, et al. Chromosome transfer induced aneuploidy results in complex dysregulation of the cellular transcriptome in immortalized and cancer cells. Cancer Res. 2004;64(19):6941–6949.

[6] Zack TI, Schumacher SE, Carter SL, Cherniack AD, Saksena G, Tabak B, et al. Pan-cancer patterns of somatic copy number alteration. Nature Genetics. 2013;45:1134–1134.

[7] Kallioniemi A, Kallioniemi O, Sudar D, Rutovitz D, Gray J, Waldman F, et al. Comparative genomic hybridization for molecular cytogenetic analysis of solid tumors. Science. 1992;5083(258):818–821.

[8] Joos S, Scherthan H, Speicher M, Schlegel J, Cremer T, Lichter P. Detection of amplified DNA sequences by reverse chromosome painting using genomic tumor DNA as probe. Hum Genet. 1993;90(6):584–589.

[9] Bentz M, Plesch A, Stilgenbauer S, Dohner H, Lichter P. Minimal sizes of deletions detected by comparative genomic hybridization. Genes Chromosomes Cancer. 1998;2(21):172–175.

[10] Solinas-Toldo S, Lampel S, Stilgenbauer S, Nickolenko J, Benner A, Dohner H, et al. Matrix-based comparative genomic hybridization: biochips to screen for genomic imbalances. Genes Chromosomes Cancer. 1997;4(20):399–407.

[11] Pinkel D, Segraves R, Sudar D, Clark S, Poole I, Kowbel D, et al. High resolution analysis of DNA copy number variation using comparative genomic hybridization to microarrays. Nat Genet. 1998;2(20):207–211.

[12] Wang D, Fan J, Siao C, Berno A, Young P, Sapolsky R, et al. Large-scale identification, mapping, and genotyping of single-nucleotide polymorphisms in the human genome. Science. 1998;280(5366):1077–1082.

[13] Zhao X, Li C, Paez J, Chin K, Jänne P, Chen T, et al. An integrated view of copy number and allelic alterations in the cancer genome using single nucleotide polymorphism arrays. Cancer Res. 2004;64(9):3060–3071.

[14] Zare F, Dow M, Monteleone N, Hosny A, Nabavi S. An evaluation of copy number variation detection tools for cancer using whole exome sequencing data. BMC Bioinformatics. 2017;18(1):286.

[15] Li S, Dou X, Gao R, Ge X, Qian M, Wan L. A remark on copy number variation detection methods. PLOS ONE. 2018;13(4):e0196226.

[16] Zhang L, Bai W, Yuan N, Du Z. Comprehensively benchmarking applications for detecting copy number variation. PLOS Computational Biology. 2019;15(5):e1007069.

[17] Macintyre G, Ylstra B, Brenton J. Sequencing Structural Variants in Cancer for Precision Therapeutics. Trends Genet. 2016;32(9):530–542.

[18] Ciriello G, Miller ML, Aksoy BA, Senbabaoglu Y, Schultz N, Sander C. Emerging landscape of oncogenic signatures across human cancers. Nature Genetics. 2013;45(10):1127–1133.

[19] Stephens PJ, Tarpey PS, Davies H, Van Loo P, Greenman C, Wedge DC, et al. The landscape of cancer genes and mutational processes in breast cancer. Nature. 2012;486(7403):400–404.

[20] Zhao S, Choi M, Overton JD, Bellone S, Roque DM, Cocco E, et al. Landscape of somatic single-nucleotide and copy-number mutations in uterine serous carcinoma. Proceedings of the National Academy of Sciences Proc Natl Acad Sci USA. 2013;110(8):2916.

[21] Grasso CS, Wu YM, Robinson DR, Cao X, Dhanasekaran SM, Khan AP, et al. The mutational landscape of lethal castration-resistant prostate cancer. Nature. 2012;487(7406):239–243.

[22] Wang K, Lim HY, Shi S, Lee J, Deng S, Xie T, et al. Genomic landscape of copy number aberrations enables the identification of oncogenic drivers in hepatocellular carcinoma. Hepatology. 2013;58(2):706–717.

[23] Juhlin CC, Goh G, Healy JM, Fonseca AL, Scholl UI, Stenman A, et al. Whole-Exome Sequencing Characterizes the Landscape of Somatic Mutations and Copy Number Alterations in Adrenocortical Carcinoma. The Journal of Clinical Endocrinology & Metabolism J Clin Endocrinol Metab. 2015;100(3):E493–E502.

[24] Hou JP, Ma J. DawnRank: discovering personalized driver genes in cancer. Genome Medicine. 2014;6(7):56.

[25] Conrad DF, Pinto D, Redon R, Feuk L, Gokcumen O, Zhang Y, et al. Origins and functional impact of copy number variation in the human genome. Nature. 2010;464(7289):704–712.

[26] Völker M, Backström N, Skinner BM, Langley EJ, Bunzey SK, Ellegren H, et al. Copy number variation, chromosome rearrangement, and their association with recombination during avian evolution. Genome Res Genome research. 2010;20(4):503–511.

[27] Chen L, Zhou W, Zhang C, Lupski JR, Jin L, Zhang F. CNV instability associated with DNA replication dynamics: evidence for replicative mechanisms in CNV mutagenesis. Human Molecular Genetics Hum Mol Genet. 2014;24(6):1574–1583.

[28] Mishra S, Whetstine JR. Different Facets of Copy Number Changes: Permanent, Transient, and Adaptive. Mol Cell Biol Molecular and cellular biology. 2016;36(7):1050–1063.

[29] Phallen J, Sausen M, Adleff V, Leal A, Hruban C, White J, et al. Direct detection of early-stage cancers using circulating tumor DNA. Science Translational Medicine. 2017;9(403):eaan2415.

[30] Li W, Zhang X, Lu X, You L, Song Y, Luo Z, et al. 5-Hydroxymethylcytosine signatures in circulating cell-free DNA as diagnostic biomarkers for human cancers. Cell Research. 2017;27(10):1243–1257.

[31] Pathak AK, Bhutani M, Kumar S, Mohan A, Guleria R. Circulating Cell-Free DNA in Plasma/Serum of Lung Cancer Patients as a Potential Screening and Prognostic Tool. Clinical Chemistry Clin Chem. 2006;52(10):1833–1842.

[32] Panagopoulou M, Karaglani M, Balgkouranidou I, Biziota E, Koukaki T, Karamitrousis E, et al. Circulating cell-free DNA in breast cancer: size profiling, levels, and methylation patterns lead to prognostic and predictive classifiers. Oncogene. 2019;38(18):3387–3401.

[33] Huang CC, Du M, Wang L. Bioinformatics Analysis for Circulating Cell-Free DNA in Cancer. Cancers. 2019;11(6).

[34] Heitzer E, Ulz P, Belic J, Gutschi S, Quehenberger F, Fischereder K, et al. Tumor-associated copy number changes in the circulation of patients with prostate cancer identified through wholegenome sequencing. Genome Medicine. 2013;5(4):30.

[35] Dawson SJ, Tsui DWY, Murtaza M, Biggs H, Rueda OM, Chin SF, et al. Analysis of Circulating Tumor DNA to Monitor Metastatic Breast Cancer. New England Journal of Medicine N Engl J Med. 2013;368(13):1199–1209.

[36] Leary RJ, Sausen M, Kinde I, Papadopoulos N, Carpten JD, Craig D, et al. Detection of Chromosomal Alterations in the Circulation of Cancer Patients with Whole-Genome Sequencing. Science Translational Medicine. 2012;4(162):162ra154.

[37] Gao B, Baudis M. Minimum error calibration and normalization for genomic copy number analysis. Genomics. 2020;112(5):3331–3341.

[38] Cai H, Kumar N, Ai N, Gupta S, Rath P, Baudis M. Progenetix: 12 years of oncogenomic data curation. Nucleic Acids Research Nucleic Acids Res. 2013;42(D1):D1055–D1062.

[39] Hutter C, Zenklusen JC. The Cancer Genome Atlas: Creating Lasting Value beyond Its Data. Cell. 2018;173(2):283–285.

[40] Campbell PJ, Getz G, Korbel JO, Stuart JM, Jennings JL, Stein LD, et al. Pan-cancer analysis of whole genomes. Nature. 2020;578(7793):82–93.

[41] Kuhn RM, Haussler D, Kent WJ. The UCSC genome browser and associated tools. Briefings in Bioinformatics. 2013;14(2):144–161.

[42] Gao B, Huang Q, Baudis M. segment liftover: a Python tool to convert segments between genome assemblies. F1000Res F1000Research. 2018;7:319.

[43] Hastings RJ, Bown N, Tibiletti MG, Debiec-Rychter M, Vanni R, Espinet B, et al. Guidelines for cytogenetic investigations in tumours. European Journal of Human Genetics. 2016;24(1):6–13.

[44] Krijgsman O, Carvalho B, Meijer GA, Steenbergen RDM, Ylstra B. Focal chromosomal copy number aberrations in cancer—Needles in a genome haystack. Biochimica et Biophysica Acta (BBA) - Molecular Cell Research. 2014;1843(11):2698–2704.

[45] Pei G, Hu R, Dai Y, Zhao Z, Jia P. Decoding whole-genome mutational signatures in 37 human pan-cancers by denoising sparse autoencoder neural network. Oncogene. 2020;39(27):5031–5041.

[46] Speech feature denoising and dereverberation via deep autoencoders for noisy reverberant speech recognition; 2014.

[47] Medical Image Denoising Using Convolutional Denoising Autoencoders; 2016.

[48] Cai H, Gupta S, Rath P, Ai N, Baudis M. arrayMap 2014: an updated cancer genome resource. Nucleic Acids Research Nucleic Acids Res. 2015;43(D1):D825–D830.

[49] World HO. International classification of diseases for oncology (ICD-O) – 3rd edition, 1st revision. Geneva; 2013.

[50] Mermel CH, Schumacher SE, Hill B, Meyerson ML, Getz G. GISTIC2.0 facilitates sensitive and confident localization of the targets of focal somatic copy-number alteration in human cancers. Genome Biology. 2011;12(4):R41.

[51] van Gent DC, Hoeijmakers JHJ, Kanaar R. Chromosomal stability and the DNA double-stranded break connection. Nature Reviews Genetics. 2001;2(3):196–206.

[52] Hastings PJ, Lupski JR, Rosenberg SM, Ira G. Mechanisms of change in gene copy number. Nat Rev Genet Nature reviews Genetics. 2009;10(8):551–564.

[53] Pihan GA, Purohit A, Wallace J, Knecht H, Woda B, Quesenberry P, et al. Centrosome Defects and Genetic Instability in Malignant Tumors. Cancer Research Cancer Res. 1998;58(17):3974.

[54] Bignell GR, Greenman CD, Davies H, Butler AP, Edkins S, Andrews JM, et al. Signatures of mutation and selection in the cancer genome. Nature. 2010;463(7283):893–898.

[55] Daugherty LC, Seal RL, Wright MW, Bruford EA. Gene family matters: expanding the HGNC resource. Human Genomics. 2012;6(1):4.

[56] Nishida K, Tamura A, Nakazawa N, Ueda Y, Abe T, Matsuda F, et al. The Ig Heavy Chain Gene Is Frequently Involved in Chromosomal Translocations in Multiple Myeloma and Plasma Cell Leukemia as Detected by In Situ Hybridization. Blood. 1997;90(2):526–534.

[57] Othman MAK, Grygalewicz B, Pienkowska-Grela B, Rygier J, Ejduk A, Rincic M, et al. A novel IGH@ gene rearrangement associated with CDKN2A/B deletion in young adult B-cell acute lymphoblastic leukemia. Oncol Lett Oncology letters. 2016;11(3):2117–2122.

[58] Babbage G, Ottensmeier CH, Blaydes J, Stevenson FK, Sahota SS. Immunoglobulin Heavy Chain Locus Events and Expression of Activation-Induced Cytidine Deaminase in Epithelial Breast Cancer Cell Lines. Cancer Research Cancer Res. 2006;66(8):3996.

[59] Hu D, Zheng H, Liu H, Li M, Ren W, Liao W, et al. Immunoglobulin expression and its biological significance in cancer cells. Cell Mol Immunol Cellular & molecular immunology. 2008;5:319–324.

[60] Lee G, Ge B. Cancer cell expressions of immunoglobulin heavy chains with unique carbohydrate-associated biomarker. Cancer Biomarkers. 2009;5:177–188.

[61] Larsson C, Ehinger A, Winslow S, Leandersson K, Klintman M, Dahl L, et al. Prognostic implications of the expression levels of different immunoglobulin heavy chain-encoding RNAs in early breast cancer. npj Breast Cancer. 2020;6(1):28.

[62] Hu D, Duan Z, Li M, Jiang Y, Liu H, Zheng H, et al. Heterogeneity of aberrant immunoglobulin expression in cancer cells. Cellular & Molecular Immunology. 2011;8(6):479–485.

[63] Weber L, Maßberg D, Becker C, Altmüller J, Ubrig B, Bonatz G, et al. Olfactory Receptors as Biomarkers in Human Breast Carcinoma Tissues. Front Oncol Frontiers in oncology. 2018;8:33.

[64] Masjedi S, Zwiebel LJ, Giorgio TD. Olfactory receptor gene abundance in invasive breast carcinoma. Scientific Reports. 2019;9(1):13736.

[65] Li M, Wang X, Ma RR, Shi DB, Wang YW, Li XM, et al. The Olfactory Receptor Family 2, Sub-family T, Member 6 (OR2T6) Is Involved in Breast Cancer Progression via Initiating Epithelial-Mesenchymal Transition and MAPK/ERK Pathway. Frontiers in Oncology. 2019;9:1210.

[66] Ranzani M, Iyer V, Ibarra-Soria X, Del Castillo Velasco-Herrera M, Garnett M, Logan D, et al. Revisiting olfactory receptors as putative drivers of cancer [version 1; peer review: 2 approved]. Wellcome Open Research. 2017;2(9).

[67] Jaworska AM, Wlodarczyk NA, Mackiewicz A, Czerwinska P. The role of TRIM family proteins in the regulation of cancer stem cell self-renewal. STEM CELLS. 2020;38(2):165–173.

[68] Mandell MA, Saha B, Thompson TA. The Tripartite Nexus: Autophagy, Cancer, and Tripartite Motif-Containing Protein Family Members. Frontiers in Pharmacology. 2020;11:308.

[69] Hatakeyama S. TRIM proteins and cancer. Nature Reviews Cancer. 2011;11:792–804.

[70] Li B, Huang Q, Wei GH. The Role of HOX Transcription Factors in Cancer Predisposition and Progression. Cancers (Basel) Cancers. 2019;11(4):528.

[71] Shah N, Sukumar S. The Hox genes and their roles in oncogenesis. Nature Reviews Cancer. 2010;10(5):361–371.

[72] Bhatlekar S, Fields JZ, Boman BM. HOX genes and their role in the development of human cancers. Journal of Molecular Medicine. 2014;92(8):811–823.

[73] Brotto DB, Siena II, Carvalho SdCeS, Muys BR, Goedert L, Cardoso C, et al. Contributions of HOX genes to cancer hallmarks: Enrichment pathway analysis and review. Tumor Biology Tumour Biol. 2020;42(5):1010428320918050.

[74] Sandrine GL, Juillerat-Jeanneret L. Glycosylation Pathways as Drug Targets for Cancer: Glycosidase Inhibitors. Mini-Reviews in Medicinal Chemistry. 2006;6(9):1043–1052.

[75] Bernacki RJ, Niedbala MJ, Korytnyk W. Glycosidases in cancer and invasion. Cancer and Metastasis Reviews. 1985;4(1):81–101.

[76] Wu Y, Chen X, Wang S, Wang S. Advances in the relationship between glycosyltransferases and multidrug resistance in cancer. Clinica Chimica Acta. 2019;495:417–421.

[77] Andergassen U, Liesche F, Kölbl AC, Ilmer M, Hutter S, Friese K, et al. Glycosyltransferases as Markers for Early Tumorigenesis. BioMed Research International. 2015;2015:792672.

[78] Strouhalova K, Přechová M, Gandalovičová A, Brábek J, Gregor M, Rosel D. Vimentin Intermediate Filaments as Potential Target for Cancer Treatment. Cancers. 2020;12(1).

[79] Sharma P, Alsharif S, Fallatah A, Chung BM. Intermediate Filaments as Effectors of Cancer Development and Metastasis: A Focus on Keratins, Vimentin, and Nestin. Cells. 2019;8(5):497.

[80] Holle AW, Kalafat M, Ramos AS, Seufferlein T, Kemkemer R, Spatz JP. Intermediate filament reorganization dynamically influences cancer cell alignment and migration. Scientific Reports. 2017;7(1):45152.

[81] Munro D, Ghersi D, Singh M. Two critical positions in zinc finger domains are heavily mutated in three human cancer types. PLOS Computational Biology. 2018;14(6):e1006290.

[82] Jen J, Wang YC. Zinc finger proteins in cancer progression. J Biomed Sci Journal of biomedical science. 2016;23(1):53.

[83] Shao X, Lv N, Liao J, Long J, Xue R, Ai N, et al. Copy number variation is highly correlated with differential gene expression: a pan-cancer study. BMC Medical Genetics. 2019;20(1):175.

[84] Bhattacharya A, Bense RD, Urzuá-Traslaviña CG, de Vries EGE, van Vugt MATM, Fehrmann RSN. Transcriptional effects of copy number alterations in a large set of human cancers. Nature Communications. 2020;11(1):715.

[85] Xie T, d’ Ario G, Lamb JR, Martin E, Wang K, Tejpar S, et al. A Comprehensive Characterization of Genome-Wide Copy Number Aberrations in Colorectal Cancer Reveals Novel Oncogenes and Patterns of Alterations. PLOS ONE. 2012;7:e42001.

[86] Kwong LN, Chin L. Chromosome 10, frequently lost in human melanoma, encodes multiple tumor-suppressive functions. Cancer Res Cancer research. 2014;74(6):1814–1821.

[87] Bax DA, Mackay A, Little SE, Carvalho D, Viana-Pereira M, Tamber N, et al. A Distinct Spectrum of Copy Number Aberrations in Pediatric High-Grade Gliomas. Clinical Cancer Research Clin Cancer Res. 2010;16(13):3368.

[88] Castells A, Gusella JF, Ramesh V, Rustgi AK. A Region of Deletion on Chromosome 22q13 Is Common to Human Breast and Colorectal Cancers. Cancer Research Cancer Res. 2000;60(11):2836.

[89] Morikawa A, Hayashi T, Kobayashi M, Kato Y, Shirahige K, Itoh T, et al. Somatic copy number alterations have prognostic impact in patients with ovarian clear cell carcinoma. Oncol Rep Oncology Reports. 2018;40(1):309–318.

[90] McNulty SN, Cottrell CE, Vigh-Conrad KA, Carter JH, Heusel JW, Ansstas G, et al. Beyond sequence variation: assessment of copy number variation in adult glioblastoma through targeted tumor somatic profiling. Human Pathology. 2019;86:170–181.

[91] Janjigian YY, Tang LH, Coit DG, Kelsen DP, Francone TD, Weiser, et al. MET Expression and Amplification in Patients with Localized Gastric Cancer. Cancer Epidemiology Biomarkers & Prevention Cancer Epidemiol Biomarkers Prev. 2011;20(5):1021.

[92] Trombetta D, Magnusson L, von Steyern FV, Hornick JL, Fletcher CDM, Mertens F. Translocation t(7;19)(q22;q13)—a recurrent chromosome aberration in pseudomyogenic hemangioendothelioma. Cancer Genetics. 2011;204(4):211–215.

[93] Taoudi Benchekroun M, Saintigny P, Thomas SM, El-Naggar AK, Papadimitrakopoulou V, Ren H, et al. Epidermal Growth Factor Receptor Expression and Gene Copy Number in the Risk of Oral Cancer. Cancer Prevention Research Cancer Prev Res (Phila). 2010;3(7):800.

[94] Kang U Ji. Characterization of amplification patterns and target genes on the short arm of chromosome 7 in early-stage lung adenocarcinoma. Mol Med Rep Molecular Medicine Reports. 2013;8(5):1373–1378.

[95] Moelans CB, de Weger RA, Monsuur HN, Vijzelaar R, van Diest PJ. Molecular profiling of invasive breast cancer by multiplex ligation-dependent probe amplification-based copy number analysis of tumor suppressor and oncogenes. Modern Pathology. 2010;23(7):1029–1039.

[96] El Gammal AT, Brüchmann M, Zustin J, Isbarn H, Hellwinkel OJC, Köllermann J, et al. Chromosome Deletions and Gains are Associated with Tumor Progression and Poor Prognosis in Prostate Cancer. Clinical Cancer Research Clin Cancer Res. 2010;16(1):56.

[97] Weiss J, Sos ML, Seidel D, Peifer M, Zander T, Heuckmann JM, et al. Frequent and Focal FGFR Amplification Associates with Therapeutically Tractable FGFR1 Dependency in Squamous Cell Lung Cancer. Science Translational Medicine. 2010;2(62):62ra93.

[98] Milella M, Falcone I, Conciatori F, Cesta Incani U, Del Curatolo A, Inzerilli N, et al. PTEN: Multiple Functions in Human Malignant Tumors. Front Oncol Frontiers in oncology. 2015;5:24.

[99] Fusco N, Sajjadi E, Venetis K, Gaudioso G, Lopez G, Corti C, et al. PTEN Alterations and Their Role in Cancer Management: Are We Making Headway on Precision Medicine. Genes. 2020;11(7).

[100] Parsons DW, Li M, Zhang X, Jones S, Leary RJ, Lin JCH, et al. The Genetic Landscape of the Childhood Cancer Medulloblastoma. Science. 2011;331(6016):435.

[101] Cross NA, Rennie IG, Murray AK, Sisley K. The identification of chromosome abnormalities associated with the invasive phenotype of uveal melanoma in vitro. Clinical & Experimental Metastasis. 2005;22(2):107–113.

[102] Cohen A, Sato M, Aldape K, Mason CC, Alfaro-Munoz K, Heathcock L, et al. DNA copy number analysis of Grade II–III and Grade IV gliomas reveals differences in molecular ontogeny including chromothripsis associated with IDH mutation status. Acta Neuropathologica Communications. 2015;3(1):34.

[103] Boots-Sprenger SHE, Sijben A, Rijntjes J, Tops BBJ, Idema AJ, Rivera AL, et al. Significance of complete 1p/19q co-deletion, IDH1 mutation and MGMT promoter methylation in gliomas: use with caution. Modern Pathology. 2013;26(7):922–929.

[104] Mathieu V, Pirker C, Schmidt WM, Spiegl-Kreinecker S, Lötsch D, Heffeter P, et al. Aggressiveness of human melanoma xenograft models is promoted by aneuploidy-driven gene expression deregulation. Oncotarget. 2012;3(4):399–413.

[105] Park AK, Lee JY, Cheong H, Ramaswamy V, Park SH, Kool M, et al. Subgroup-specific prognostic signaling and metabolic pathways in pediatric medulloblastoma. BMC Cancer. 2019;19(1):571.

[106] Azizi E, Friedman J, Pavlotsky F, Iscovich J, Bornstein A, Shafir R, et al. Familial cutaneous malignant melanoma and tumors of the nervous system. Cancer. 1995;76(9):1571–1578.

[107] Scarbrough PM, Akushevich I, Wrensch M, Il’yasova D. Exploring the association between melanoma and glioma risks. Ann Epidemiol Annals of epidemiology. 2014;24(6):469–474.

[108] Desai AS, Grossman SA. Association of melanoma with glioblastoma multiforme. Journal of Clinical Oncology JCO. 2008;26(15 suppl):2082.

[109] Middleton MR, Grob JJ, Aaronson N, Fierlbeck G, Tilgen W, Seiter S, et al. Randomized Phase III Study of Temozolomide Versus Dacarbazine in the Treatment of Patients With Advanced Metastatic Malignant Melanoma. Journal of Clinical Oncology JCO. 2000;18(1):158.

[110] Powell DR, O’Brien JH, Ford HL, Artinger KB. Chapter 16 - Neural Crest Cells and Cancer: Insights into Tumor Progression Neural Crest Cells. In: Trainor PA, editor. Neural Crest Cells. Boston: Academic Press; 2014. p. 335–357.

[111] Maguire LH, Thomas AR, Goldstein AM. Tumors of the neural crest: Common themes in development and cancer. Developmental Dynamics Dev Dyn. 2015;244(3):311–322.

[112] Jiang M, Stanke J, Lahti JM. The connections between neural crest development and neuroblastoma. Curr Top Dev Biol Current topics in developmental biology. 2011;94:77–127.

[113] Kovacs D, Migliano E, Muscardin L, Silipo V, Catricalà C, Picardo M, et al. The role of Wnt/beta-catenin signaling pathway in melanoma epithelial-to-mesenchymal-like switching: evidences from patients-derived cell lines. Oncotarget. 2016;7(28):43295–43314.

[114] Sinnberg T, Levesque MP, Krochmann J, Cheng PF, Ikenberg K, Meraz-Torres F, et al. Wnt-signaling enhances neural crest migration of melanoma cells and induces an invasive phenotype. Molecular Cancer. 2018;17(1):59.

[115] Zuccarini M, Giuliani P, Ziberi S, Carluccio M, Iorio PD, Caciagli F, et al. The Role of Wnt Signal in Glioblastoma Development and Progression: A Possible New Pharmacological Target for the Therapy of This Tumor. Genes (Basel) Genes. 2018;9(2):105.

[116] Doussouki ME, Gajjar A, Chamdine O. Molecular genetics of medulloblastoma in children: diagnostic, therapeutic and prognostic implications. Future Neurology. 2019;14(1):FNL8.

[117] Gallik KL, Treffy RW, Nacke LM, Ahsan K, Rocha M, Green-Saxena A, et al. Neural crest and cancer: Divergent travelers on similar paths. Mech Dev Mechanisms of development. 2017;148:89–99.

[118] Wood-Bouwens C, Lau BT, Handy CM, Lee H, Ji HP. Single-Color Digital PCR Provides High-Performance Detection of Cancer Mutations from Circulating DNA. J Mol Diagn The Journal of molecular diagnostics: JMD. 2017;19(5):697–710.

[119] Volik S, Alcaide M, Morin RD, Collins C. Cell-free DNA (cfDNA): Clinical Significance and Utility in Cancer Shaped By Emerging Technologies. Molecular Cancer Research Mol Cancer Res. 2016;14(10):898.

[120] Chin RI, Chen K, Usmani A, Chua C, Harris PK, Binkley MS, et al. Detection of Solid Tumor Molecular Residual Disease (MRD) Using Circulating Tumor DNA (ctDNA). Mol Diagn Ther Molecular diagnosis & therapy. 2019;23(3):311–331.

[121] Karim MR, Rahman A, Jares JB, Decker S, Beyan O. A snapshot neural ensemble method for cancer-type prediction based on copy number variations. Neural Computing and Applications. 2020;32(19):15281–15299.

[122] Liang Y, Wang H, Yang J, Li X, Dai C, Shao P, et al. A Deep Learning Framework to Predict Tumor Tissue-of-Origin Based on Copy Number Alteration. Frontiers in Bioengineering and Biotechnology. 2020;8:701.

[123] Zhang N, Wang M, Zhang P, Huang T. Classification of cancers based on copy number variation landscapes. Biochimica et Biophysica Acta (BBA) - General Subjects Systems Genetics - Deciphering the Complex Disease with a Systems Approach. 2016;1860(11, Part B):2750–2755.

[124] Tyanova S, Albrechtsen R, Kronqvist P, Cox J, Mann M, Geiger T. Proteomic maps of breast cancer subtypes. Nature Communications. 2016;7(1):10259.

[125] Curtis C, Shah SP, Chin SF, Turashvili G, Rueda OM, Dunning MJ, et al. The genomic and transcriptomic architecture of 2,000 breast tumours reveals novel subgroups. Nature. 2012;486(7403):346–352.

[126] Banerji S, Cibulskis K, Rangel-Escareno C, Brown KK, Carter SL, Frederick AM, et al. Sequence analysis of mutations and translocations across breast cancer subtypes. Nature. 2012;486(7403):405–409.

[127] Szalai B, Saez-Rodriguez J. Why do pathway methods work better than they should. bioRxiv. 2020;p. 2020.07.30.228296.

