## Supplementary implementations for "Signatures of Discriminative Copy Number Aberrations in 31 Cancer Subtypes"

#### Data processing

A major challenge of the study was the integration of CNV data, originating from different studies and generated with different technologies and settings. In the data processing pipeline (workflow in Figure 1 of the article), we addressed the following challenges in order to produce a uniform dataset.

(1). Data format heterogeneity: We converted all original CNV data into called copy number segments.

(2). Reference genome difference: We applied a previously developed tool (*segmentLiftover*) for copy number data conversion between different genome editions. If probe specific data was available, we lifted those to GRCh38 (hg38) then applied the *Progenetix* standard segmentation pipeline to generate copy number segmentation data; otherwise, we directly lifted the original segmentation data to GRCh38. As experiments during the development of *segmentLiftover* indicated, direct conversion using segmentation data showed more than 99% accuracy compared to the gold standard method; however, conversion at probe level data still showed a marginal advantage in accuracy at the cost of computation workload.

(3). Signal calibration: mainly due to the existence of tumor sub-clones and normal cells, the values of the normal copy number state usually are not reflected as 2 (0 in log ratio) in CNV profiles, and the value differences between two copy number states are different in every profile. One solution to the problem is to detect the actual purity and ploidy of each sample, and use the variation of the primary tumor as the calibration baseline. However, purity and ploidy estimation is an extremely challenging task requiring either raw data or extensive computation power. Neither is applicable for the size of our data nor the capacity of our resource. Therefore, we developed an alternative method (*Mecan4CNA*) to identify the values corresponding to the normal level and one copy change in the primary tumor clone for each CNV profile. Without actually solving the purity and ploidy, (*Mecan4CNA*) was able to estimate parameters, which could be used for data normalization, in an efficient and relatively accurate manner.

(4). Data normalization: after applying *Mecan4CNA*, we were able to normalize all samples by the actual value of the normal level and the one copy variation of the primary tumor, where the normal copy number state became the value of 2 and the one copy alternation of the primary clone became a value of 1 or 3 (log ratio values were used in actual data). The *Supplementary Normalization Examples* illustrates how different samples are normalized and aligned in the pipeline.

(5). Data curation: the format and detail of metadata, especially the clinical information, are usually different in each study. In our research group, a standard procedure is set up in the data updating pipeline of *Progenetix* to manually examine, correct, and transform the metadata into a uniform structure. The details of the related work and procedure are elaborated in the documentation of *Progenetix* and an upcoming article of the recent major update of *Progenetix*.

(6). Quality control: data were filtered at each of the previous steps with different focuses. (i). During genome conversion, samples were excluded if more than 20% of the data were not able to be converted. The significant loss of data may considerably reduce the representation validity of the sample to its tumor type. (ii). During the computation of *Mecan4CNA*, samples were reported and removed if they were suspected to be aneuploid by detecting the abnormal distribution of signals. The aneuploid samples have reduced accuracy in estimation and introduce extra layers of complexity in normalization, which may reduce the dataset's overall consistency and comparability. (iii). After applying *Mecan4CNA*, samples were removed if the deviation of the estimations were greater than a threshold (0.3 were used, based experiments during *Mecan4CNA* development) from the expected values. These samples usually are complex heterogeneous samples or low-quality samples, which the algorithm fails to evaluate correctly. (iv). During data curation, samples with incomplete diagnostic information were tagged and excluded from the study. (v). Finally, at the beginning of the analysis, samples were removed if the number of segments is greater than 3000. These samples may be hyper-segments or of low quality, which may introduce unintended bias to the analysis.

### Comparison of autoencoders

In this study, we compared the performance of four different autoencoders (basic, denoising, sparse and contractive autoencoders) on their abilities of input reconstruction. In total, we performed three groups of tests to evaluate the impact of different core sizes, layer depths, and autoencoder types. The autoencoders were constructed with Keras in Python and used Mean Square Error (MSE) as the loss function. In the first test group, we compared the performance on a different number of nodes of the encoding layer (32, 64, 128, 256, 512, 1024, 2048, 4056). All encoders and decoders were three layers deep. The results showed that in all types, the loss decreased with the increase of core size. However, the decrease of loss became very small after 1024 nodes. In the second test group, we compared the performance on different depths of the neural network (1 to 5 hidden layers). The results showed that although the loss decreased with the increase of depths, the increase was marginal. In the third test group, we compared the performance of different parameters for each autoencoder variant. For the denoising autoencoder, the noise factor, which was generated from a Gaussian distribution, was set to [0.005, 0.01, 0.025, 0.05, 0.075, 0.1]. For the sparse autoencoder, the sparsity was achieve through L1 regularization, and the regularization parameter $\lambda$ was set to $[10^{-2}, 10^{-3}, 10^{-4}, 10^{-5}, 10^{-6}, 10^{-7}]$. For the contractive autoencoder, parameter $\lambda$ in the contractive loss function was set to $[10^{-2}, 10^{-3}, 10^{-4}, 10^{-5}, 10^{-6}, 10^{-7}]$. Figure 1 shows the comparison of performance between different autoencoder settings. Figure 2 illustrates visual comparisons of each type's ability to reconstruction the input data. In terms of restoration accuracy, the denoising and contractive autoencoders showed the best performance. Since the recursive computation of the contractive function is significantly more expensive than applying a mask of noise, the denoising autoencoder was finally selected for its efficiency.

#### Implementation of the hybrid model

The hybrid model was implemented in Python. The autoencoder was constructed using Keras, and the LRP was archived through the iNNvestigate library. In phase one, both the encoder and decoder were three layers deep. The encoding layer had 1024 nodes. The noise mask was generated using a Gaussian distribution with 0 mean and 0.05 standard deviation. In phase two, both the encoder and decoder were three layers deep. The encoding layer had 3000 nodes. The noise mask was generated using a Gaussian distribution with 0 mean and 0.05 standard deviation. In both phases, the LRP-epsilon formula was used as the updated rule to compute an importance matrix of the original features.

#### Feature selection

After the training in each phase, the Autoencoder transformed the input features into an encoding space. Then by applying the LRP algorithm, we were able to calculate a weighting matrix of the original input against the samples. A high absolute score indicates the high importance of the feature in reconstructing the sample and distinguishing it from others. Then, the matrix was summed to compute the average weighting score of each feature. In order to determine the optimal cutoff to remove low-importance features, we trained the Autoencoder iteratively on a subset of the original features generated from increasing cutoff values. The largest cutoff before the dramatic increase of reconstruction errors was used to select features. In the cytoband phase, the features were reduced from 1,622 to 159. In the gene phase, the features were reduced from 3,029 to 927.

#### Generation of signatures

The CNA signature was a further abstraction for a specific subtype based on the selected feature genes of the entire sample space. First, all samples of the subtype were transformed into the representation of feature genes. Then, the feature matrix was normalized and followed by a summation by features. Next, the distributions of feature values, which represented the frequency of mutations, were computed. Based on the distributions (Figure 3, features in the least frequent distribution were excluded, and the remaining features were selected to form the signature of the subtype.

#### Implementation of the classifier

The multi-labels classifier was constructed in Python using *scikit-learn*. Because the number of samples is extremely imbalanced between different classes, undersampling and oversampling techniques were recruited to bring the classes into a more balanced state.

In undersampling, each time, 1700 random samples were drawn from the breast infiltrating duct carcinoma class, which consisted of over 5,700 samples. In oversampling, the Synthetic Minority Oversampling Technique (SMOTE) was used to generate new samples for minor classes. Ten variants of SMOTE were compared against their accuracy and speed on the cancer subtype dataset. Each variant was tested using 10-fold cross-validation combined with a random forest classifier. As shown in Figure 4, ProWSyn showed the best overall performance among the tested variants. The implementation of SMOTE variants was based on the *SMOTE_Variants* library.

Next, a grid search was conducted in each validation iteration to optimize the hyperparameters of the random forest classifier. The following parameters: *number of estimators, minimal samples split, maximum depth*, were searched with 5-fold cross validations on the training set. The most common best models were constructed with *n_estimators=2000, min_samples_split=3, max_depath=30, random_state=514*.

Finally, the prediction performance on cancer subtypes was computed as the average of 10 iterations of under-and-over-samplings and 10-fold cross-validations. The prediction result of organs was achieved through mapping the sample labels from sub-type to their corresponding organs (topography), where the metrics could be calculated using the mapped true labels and predicted labels. Compared with the direct approach to predict organs, where a classier is trained using organs as labels, the mapping method showed around 10% improvement in overall performance.

### Appendix

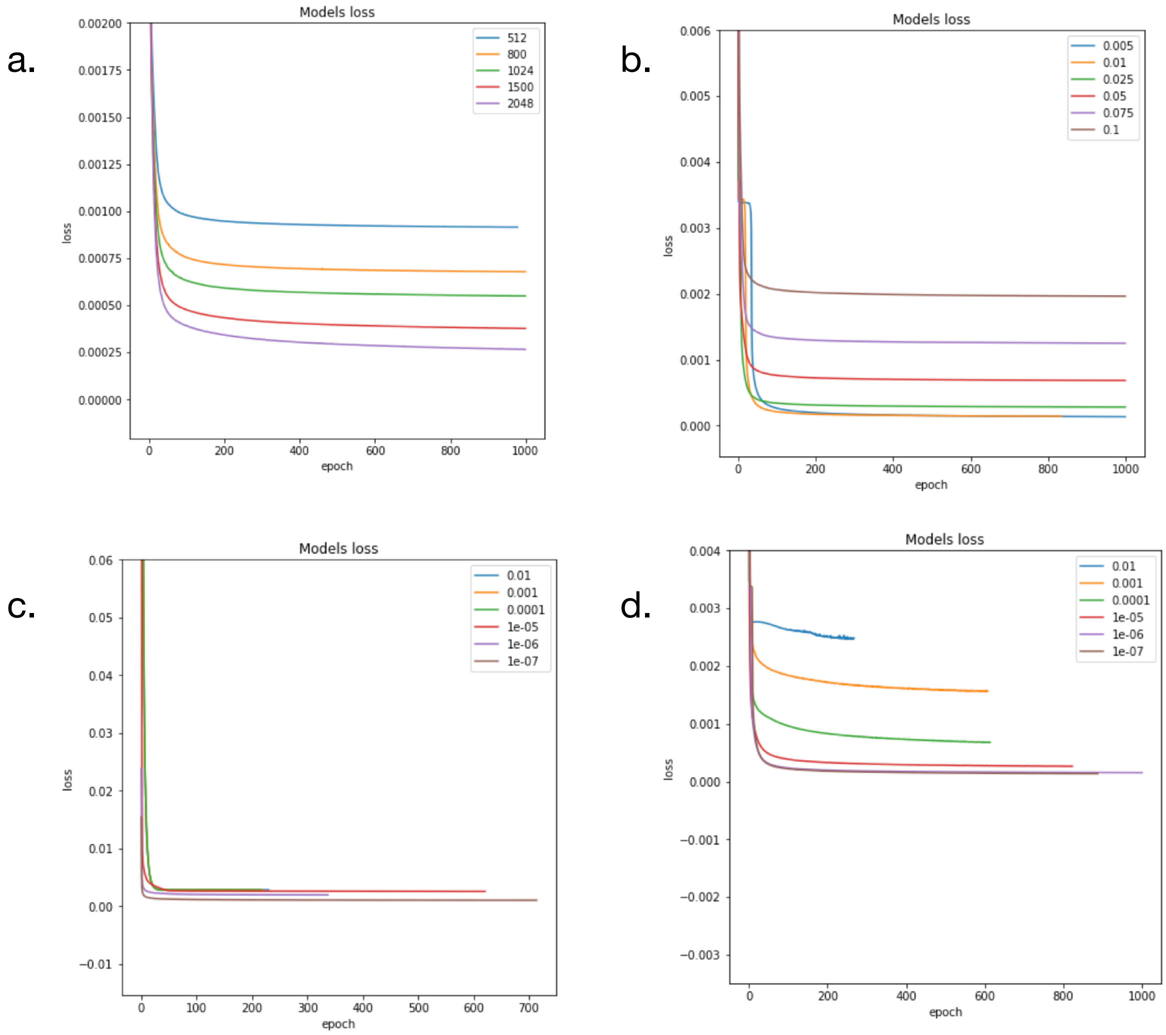


Figure 1: The comparisons of parameter settings of autoencoders. (a) is the comparison of core size. (b) is the comparison of noise factors of denoising autoencoders. (c) is the comparison of lambda of sparse autoencoders. (d) is the comparison of lambda of contractive autoencoders.

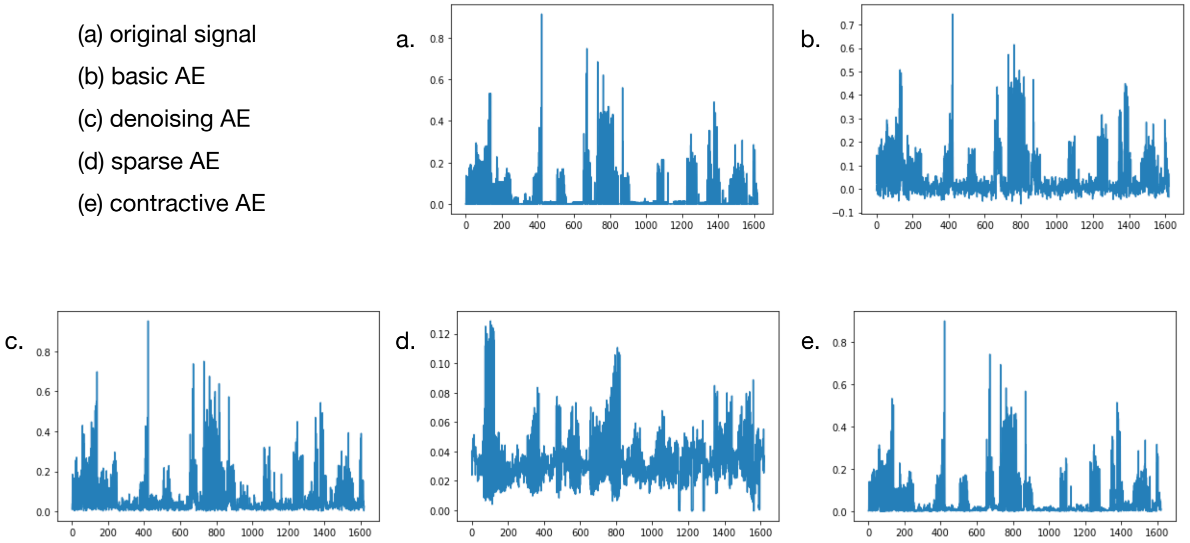


Figure 2: A visual illustration of the restoration accuracy of different autoencoders. The x-axis are features and the y-axis are normalized values. The *original signal* is the visualization of the feature space of a random sample. The rest of the figures are restoration results of different autoencoders.

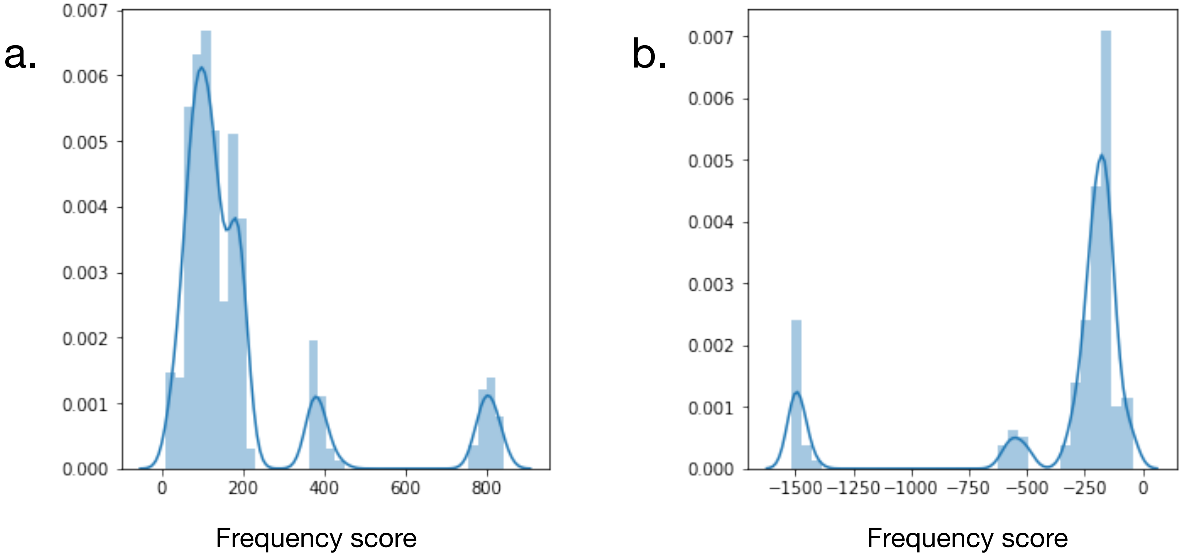


Figure 3: The distributions of the feature's frequency scores in glioma. (a) illustrates the distributions of amplifications, and (b) illustrates the distributions of deletions. A high frequency score indicates the prevalence of the feature among all samples.

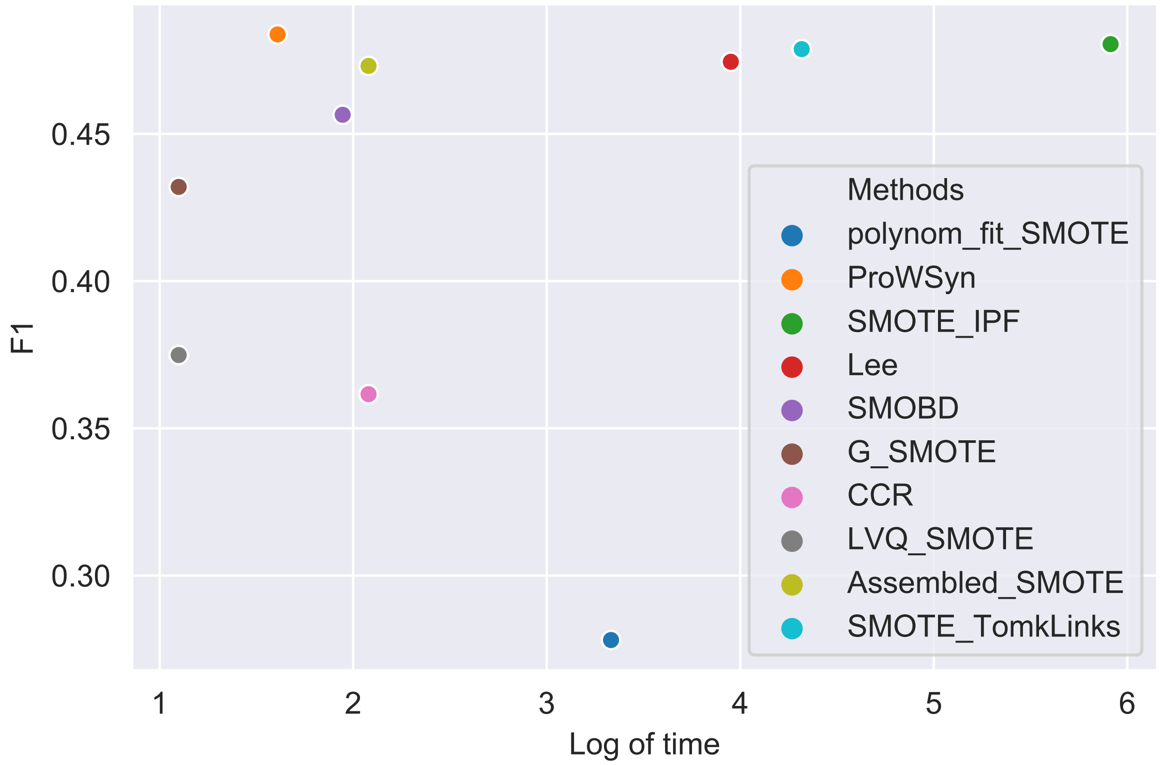


Figure 4: The comparison of ten SMOTE variants. The x-axis is the log of time in minutes.
