## Supplementary Density Plot for "Signatures of Discriminative Copy Number Aberrations in 31 Cancer Subtypes"

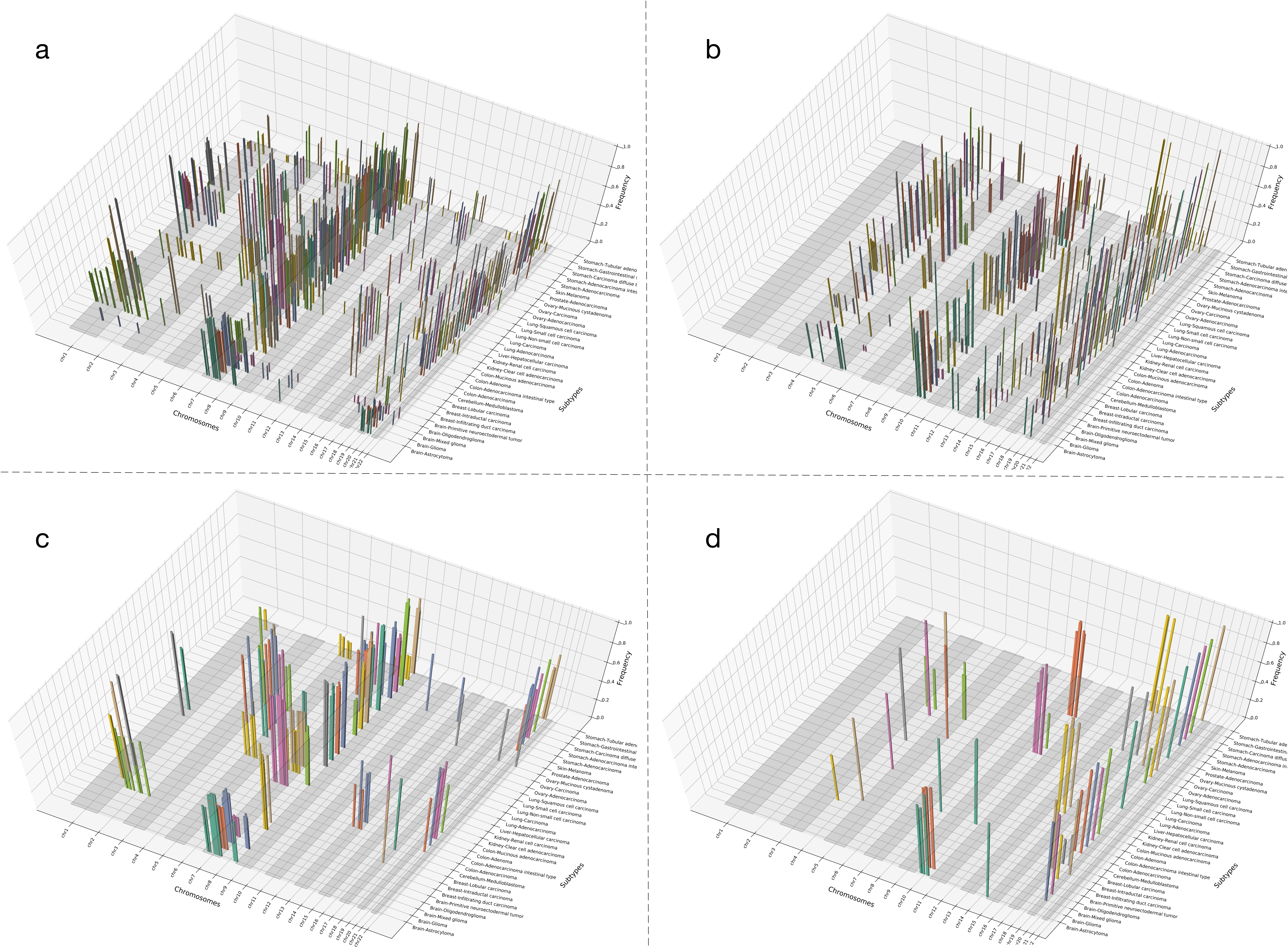


The distribution and frequency of feature genes of 31 cancer subtypes on each chromosome.
(a) shows all duplication feature genes,
(b) shows all deletion feature genes,
(c) shows only the duplication genes in signatures,
(d) shows only the deletion genes in signatures.
The signatures further reduce the complexity of feature space while maintaining the separation for each subtype.
