## Supplementary Normalization Examples for "Signatures of Discriminative Copy Number Aberrations in 31 Cancer Subtypes"

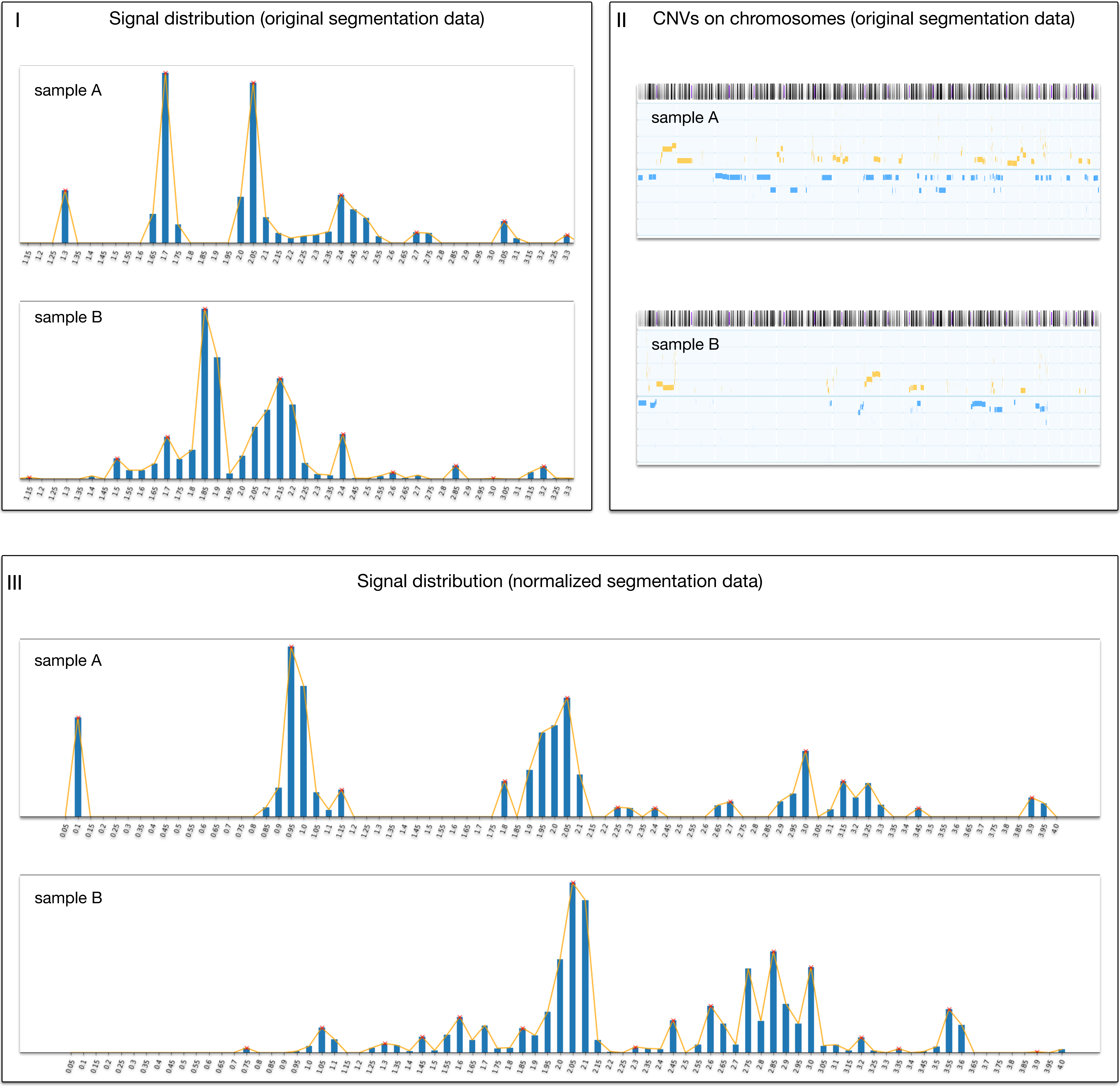


**An illustrations of CNV data normalization using Mecan4cna**

CNV data from different callsets or studies usually cannot be compared directly, because estimated copy number values have no absolute or strictly linear correlation to their corresponding DNA levels, and the extent of deviation differs between sample profiles. The figure illustrates the difference between two samples before and after normalization. Sample A is from Progenetix (callset: GSE7545, sample id: GSM182841, platform: Affymetrix Mapping 250K Nsp SNP Array). Sample B is from TCGA (project: BRCA, sample id: ae96c429-b221-4894-a45a-6aa4e8d32c71, platform: Affymetrix SNP 6 Array). Both samples are from breast cancer tumors diagnosed as infiltrating duct carcinoma, NOS (ICDO-M: 8500/3).

Fig (I) shows the CNV value distribution of the original data. The x-axis is the number of copies. The y-axis is the accumulated intensity normalized with the maximum value. Fig (II) shows the original CNV segments on chromosomes. The x-axis is chromosomes from 1 to 22. The y-axis is the CNV log2-ratio between [-2,2]. Fig (III) shows the CNV value distribution of the normalized data. The x-axis is the number of copies. The y-axis is the accumulated intensity normalized with the maximum value.
