## Supplementary Performance for "Signatures of Discriminative Copy Number Aberrations in 31 Cancer Subtypes"

Performance on individual sub-types

| Label | Precision | Recall | F1-score |
|---:|---:|---:|---:|
| Breast Intraductal carcinoma | 0.7200 | 0.7200 | 0.7200 |
| Colon Adenocarcinoma | 0.6888 | 0.7363 | 0.7118 |
| Brain Glioma | 0.7205 | 0.6784 | 0.6988 |
| Cerebellum Medulloblastoma | 0.6317 | 0.7743 | 0.6957 |
| Ovary Carcinoma | 0.6885 | 0.6535 | 0.6705 |
| Kidney Clear cell adenocarcinoma | 0.5751 | 0.7957 | 0.6677 |
| Prostate Adenocarcinoma | 0.5578 | 0.6145 | 0.5848 |
| Breast Lobular carcinoma | 0.4646 | 0.7667 | 0.5786 |
| Colon Adenoma | 0.5882 | 0.5263 | 0.5556 |
| Skin Melanoma | 0.5912 | 0.5063 | 0.5455 |
| Ovary Mucinous cystadenoma | 0.4423 | 0.6571 | 0.5287 |
| Liver Hepatocellular carcinoma | 0.4220 | 0.6577 | 0.5141 |
| Breast Infiltrating duct carcinoma | 0.5542 | 0.4412 | 0.4913 |
| Lung Squamous cell carcinoma | 0.4190 | 0.5677 | 0.4822 |
| Stomach Gastrointestinal stromal sarcoma | 0.4265 | 0.5472 | 0.4793 |
| Lung Non-small cell carcinoma | 0.4729 | 0.4710 | 0.4720 |
| Brain Oligodendroglioma | 0.4444 | 0.4848 | 0.4638 |
| Lung Adenocarcinoma | 0.5837 | 0.3736 | 0.4556 |
| Stomach Adenocarcinoma | 0.6000 | 0.3144 | 0.4126 |
| Lung Small cell carcinoma | 0.3220 | 0.4130 | 0.3619 |
| Kidney Renal cell carcinoma | 0.3261 | 0.3093 | 0.3175 |
| Brain Primitive neuroectodermal tumor | 0.3333 | 0.2903 | 0.3103 |
| Brain Astrocytoma | 0.2895 | 0.2157 | 0.2472 |
| Colon Adenocarcinoma intestinal type | 0.2222 | 0.2500 | 0.2353 |
| Stomach Tubular adenocarcinoma | 0.2222 | 0.2400 | 0.2308 |
| Ovary Adenocarcinoma | 0.2333 | 0.2121 | 0.2222 |
| Stomach Adenocarcinoma intestinal type | 0.1667 | 0.2400 | 0.1967 |
| Stomach Carcinoma diffuse type | 0.1471 | 0.2941 | 0.1961 |

| Label | Precision | Recall | F1-score |
|---|---|---|---|
| Brain Mixed glioma | 0.2286 | 0.1633 | 0.1905 |
| Lung Carcinoma | 0.2500 | 0.1220 | 0.1639 |
| Colon Mucinous adenocarcinoma | 0.1250 | 0.1579 | 0.1395 |

Performance on individual organs

| Label | Precision | Recall | F1-score |
|---:|---:|---:|---:|
| Brain | 0.7860 | 0.7174 | 0.7502 |
| Lung | 0.7524 | 0.6948 | 0.7225 |
| Colon | 0.6894 | 0.7386 | 0.7132 |
| Cerebellum | 0.6317 | 0.7743 | 0.6957 |
| Ovary | 0.6809 | 0.6661 | 0.6734 |
| Kidney | 0.5983 | 0.7606 | 0.6698 |
| Prostate | 0.5578 | 0.6145 | 0.5848 |
| Breast | 0.6000 | 0.5345 | 0.5653 |
| Skin | 0.5912 | 0.5063 | 0.5455 |
| Liver | 0.4220 | 0.6577 | 0.5141 |
| Stomach | 0.5228 | 0.4269 | 0.4700 |
