## Supplementary figures and images for "Signatures of Discriminative Copy Number Aberrations in 31 Cancer Subtypes"

### sigGenes.pdf

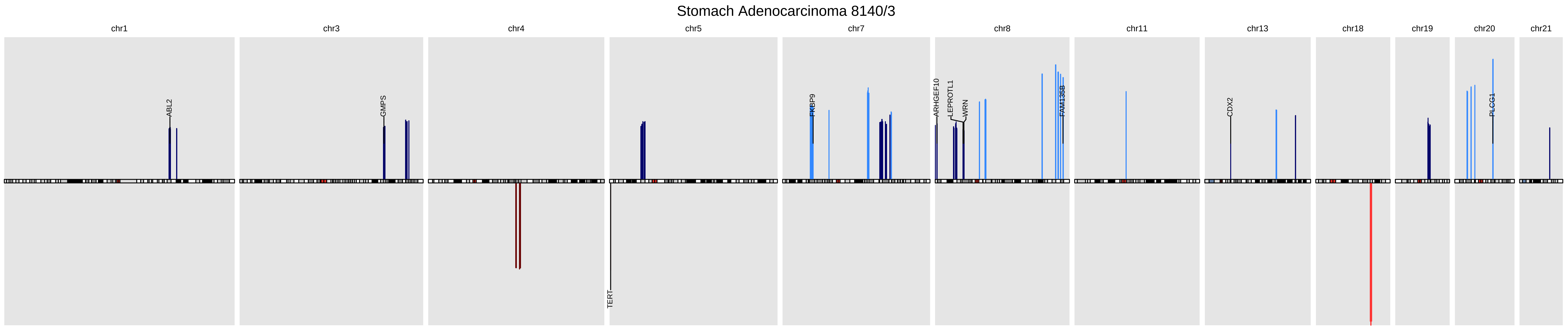

### sigGenes_full.pdf

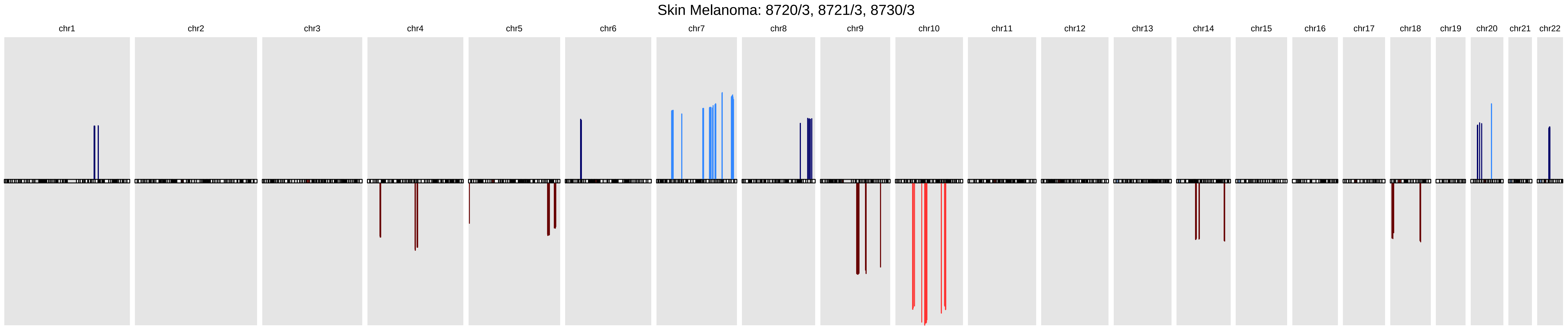

### sigGenes_full.pdf

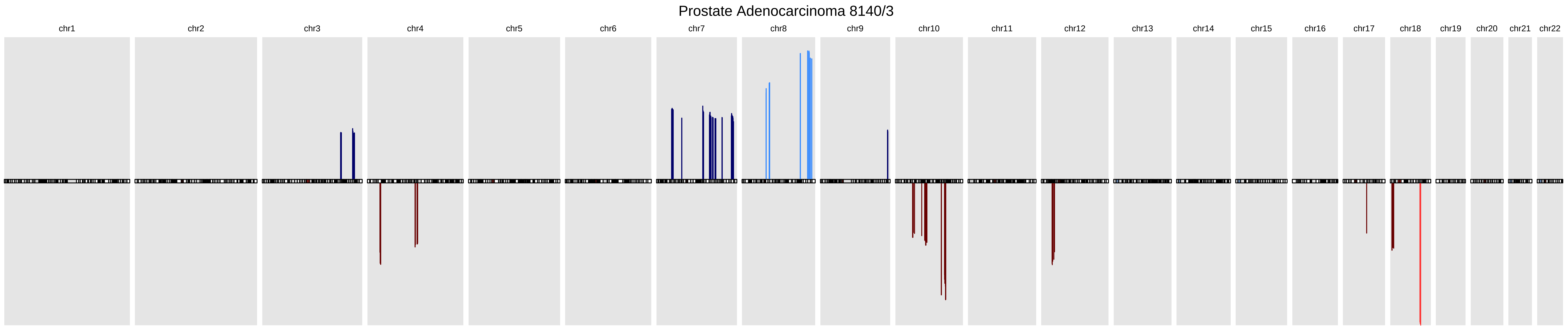

### sigGenes_full.pdf

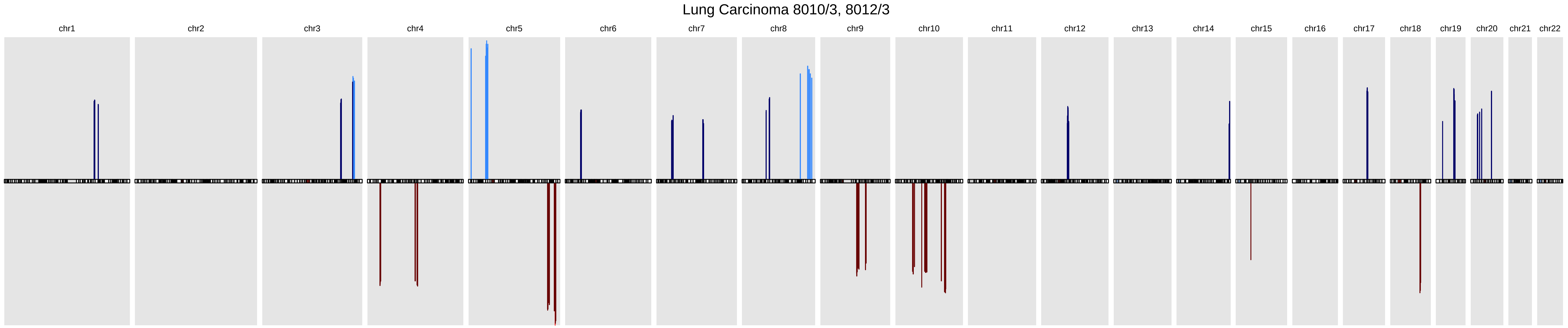

### sigGenes_full.pdf

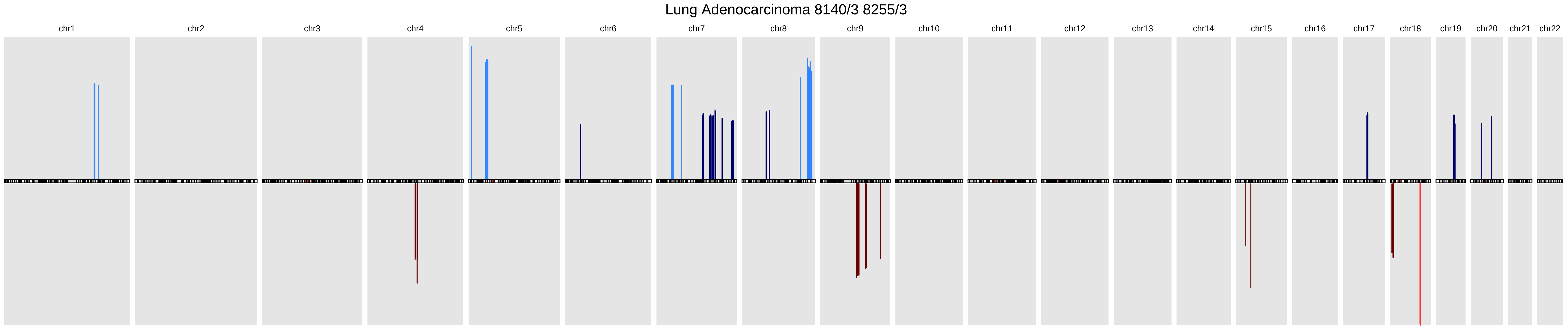

### sigGenes_full.pdf

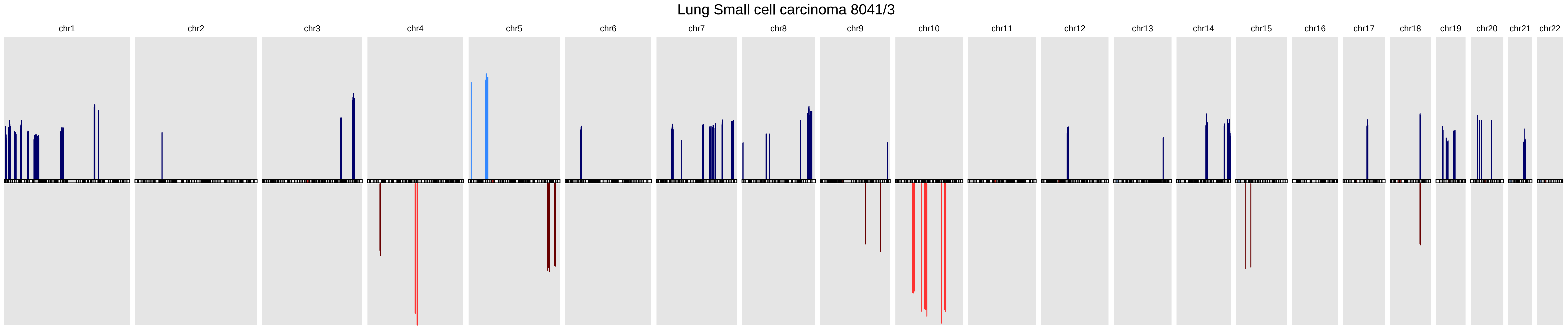

### sigGenes_full.pdf

#### Lung Squamous cell carcinoma 8070/3

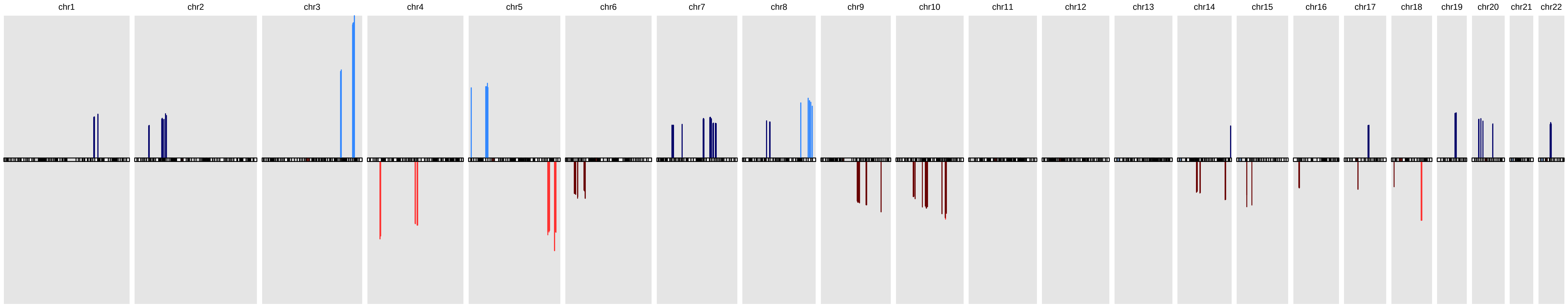

### sigGenes_full.pdf

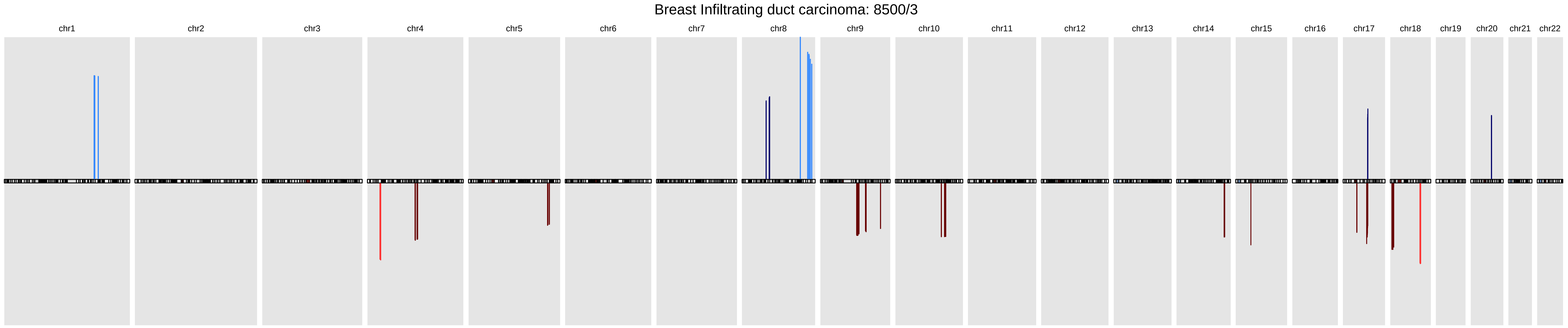

### sigGenes_full.pdf

#### Breast Intraductal carcinoma: 8500/2

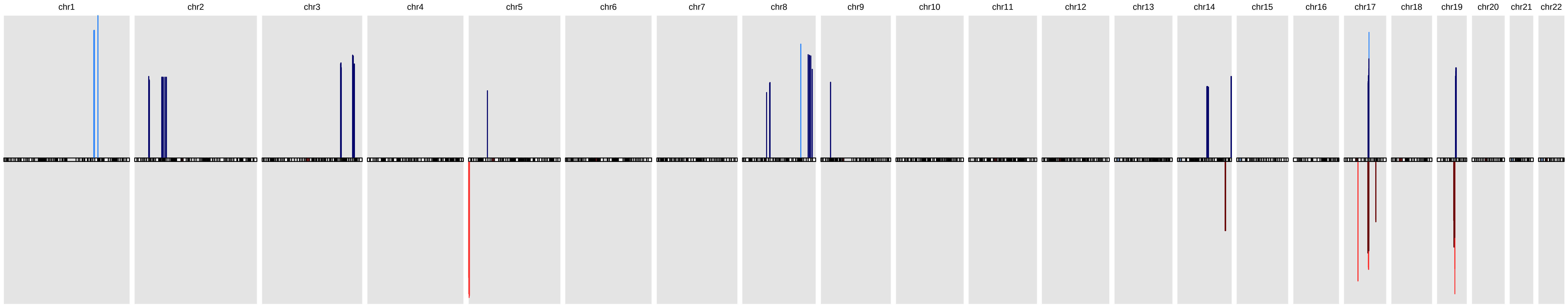

### sigGenes_full.pdf

#### Breast Lobular carcinoma: 8520/3

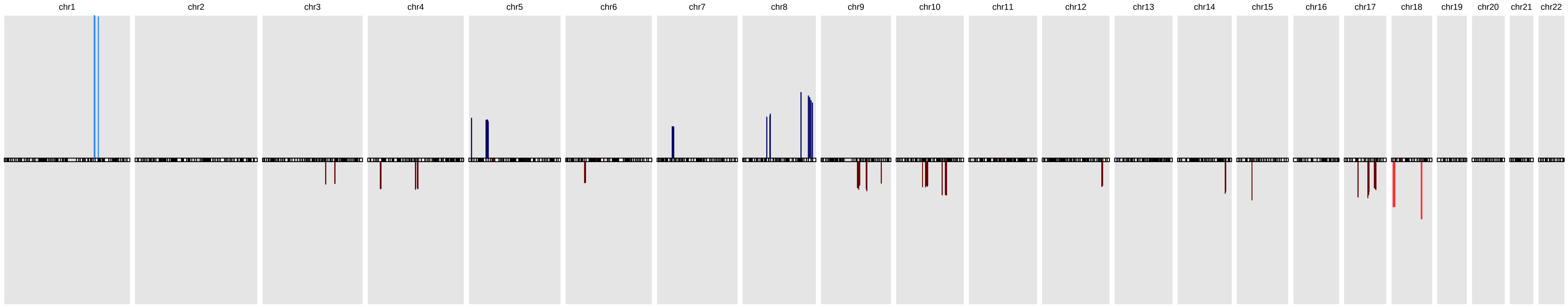

### sigGenes_full.pdf

#### Kidney Renal cell carcinoma 8312/3

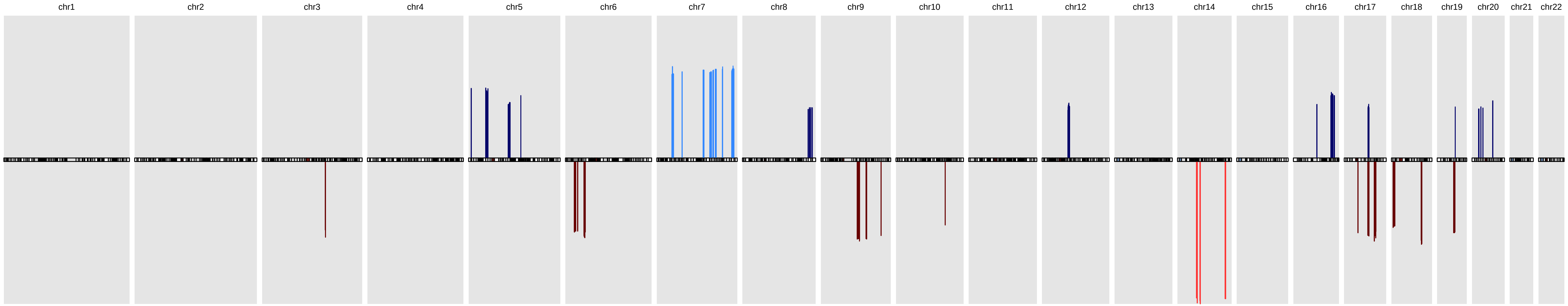

### sigGenes_full.pdf

#### Kidney Clear cell adenocarcinoma 8310/3

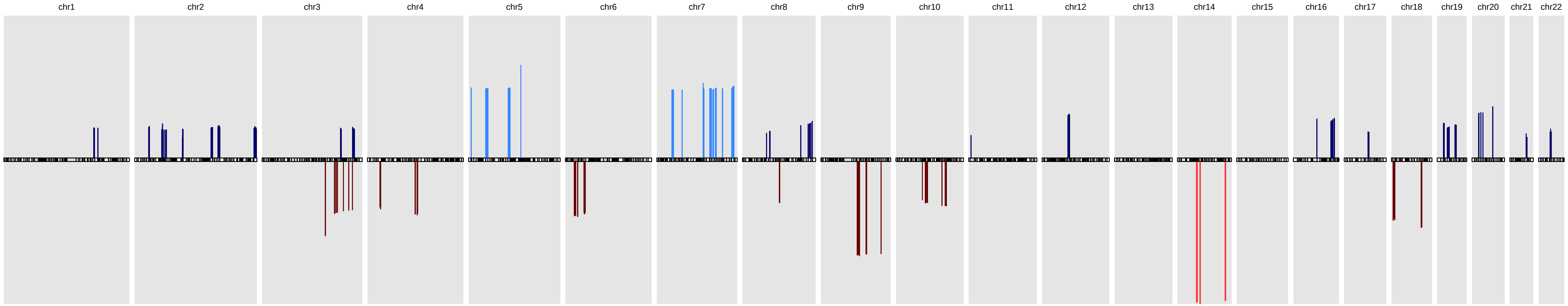

### sigGenes_full.pdf

#### Colon Adenoma: 8140/0

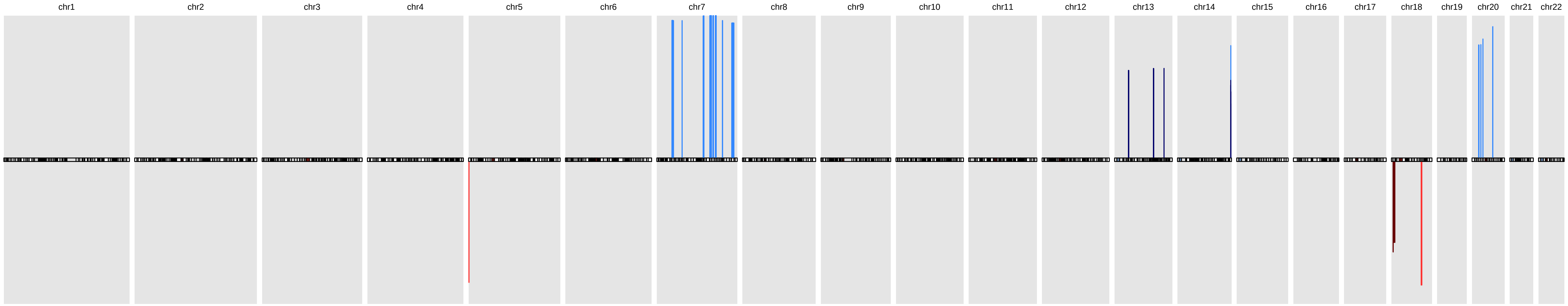

### sigGenes_full.pdf

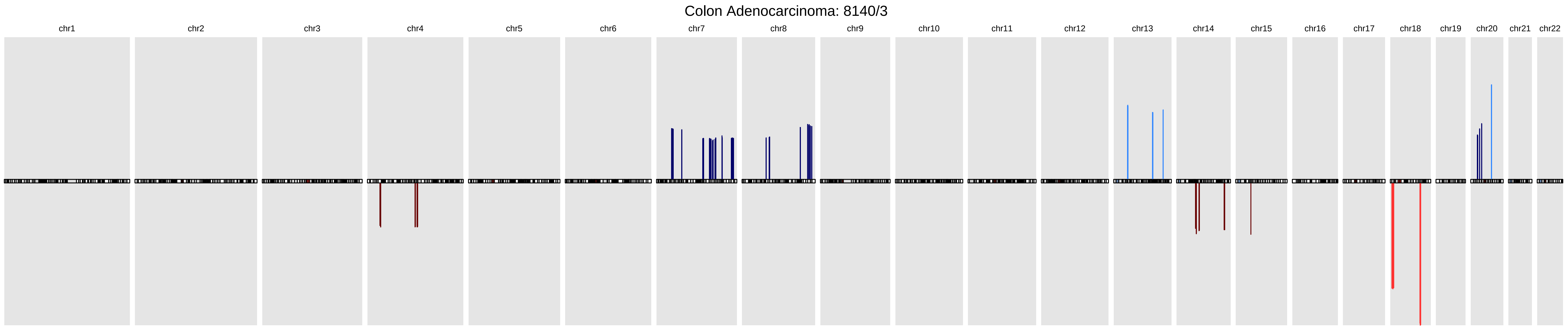

### sigGenes_full.pdf

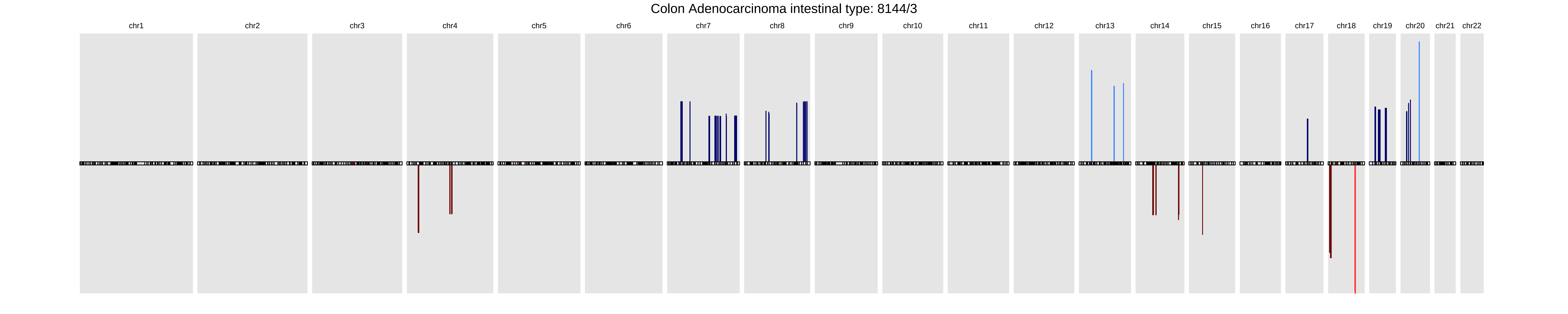

### sigGenes_full.pdf

##### Colon Mucinous adenocarcinoma: 8480/3

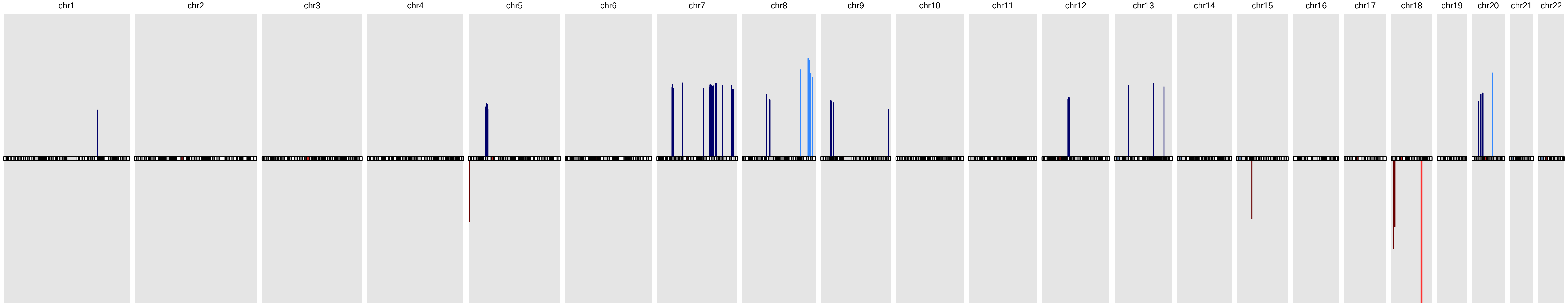

### sigGenes_full.pdf

#### Cerebellum Medulloblastoma 9470/3, 9471/3, 9474/3

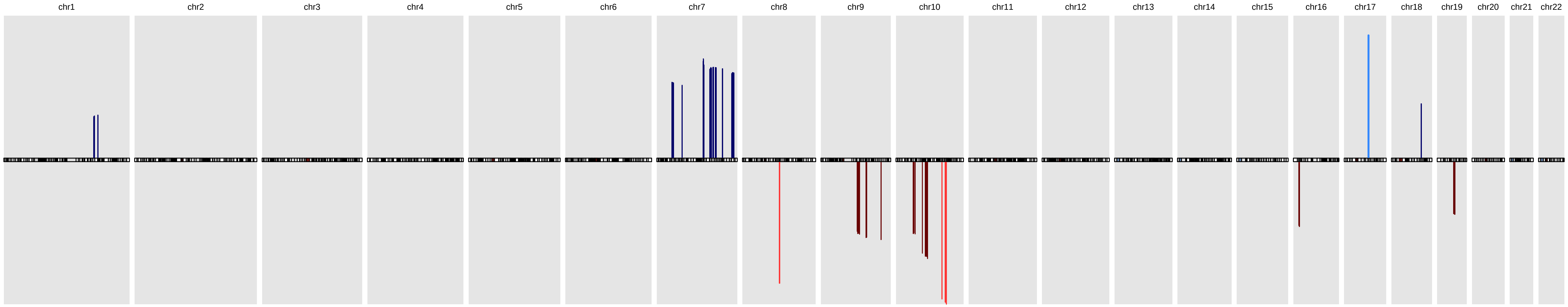

### sigGenes_full.pdf

##### Stomach Gastrointestinal stromal sarcoma 8936/3

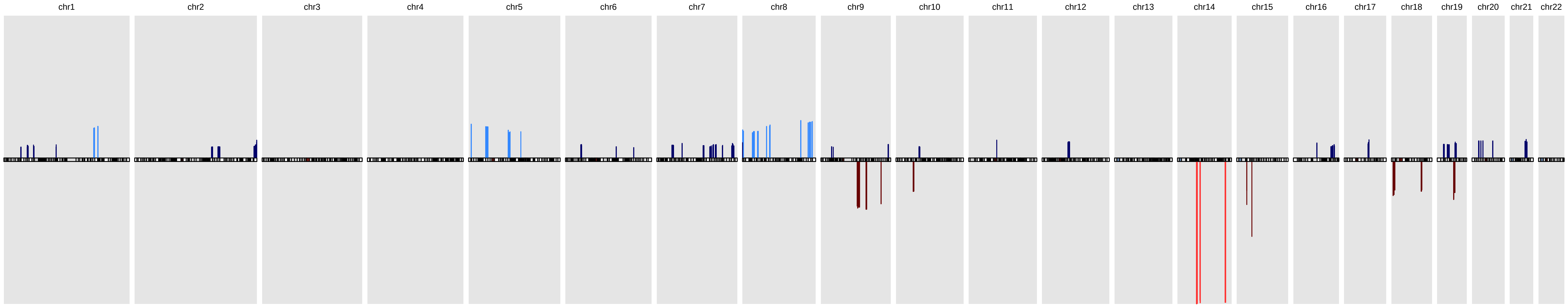

### sigGenes_full.pdf

Stomach Carcinoma diffuse type 8145/3

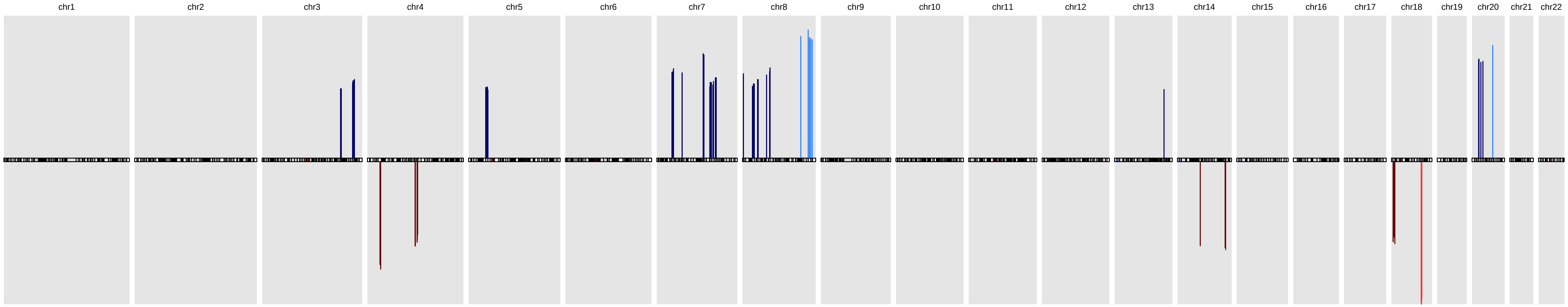

### sigGenes_full.pdf

#### Stomach Adenocarcinoma intestinal type 8144/3

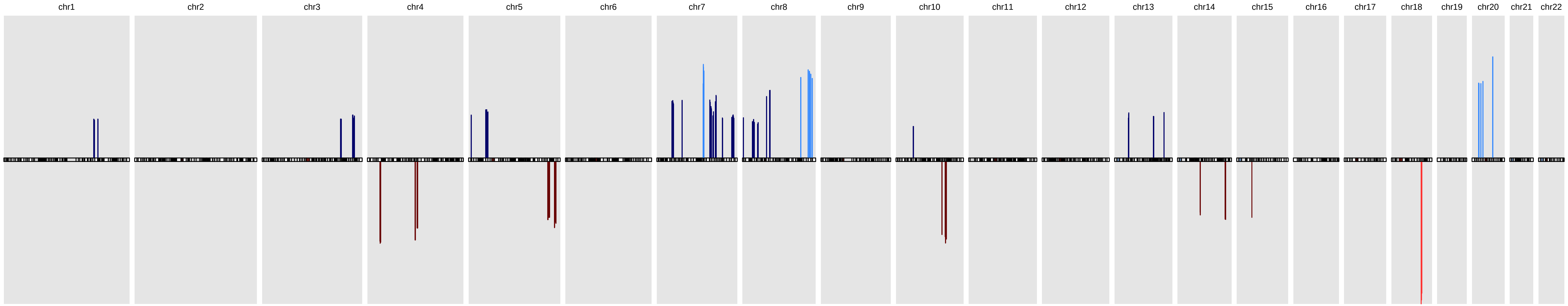

### sigGenes_full.pdf

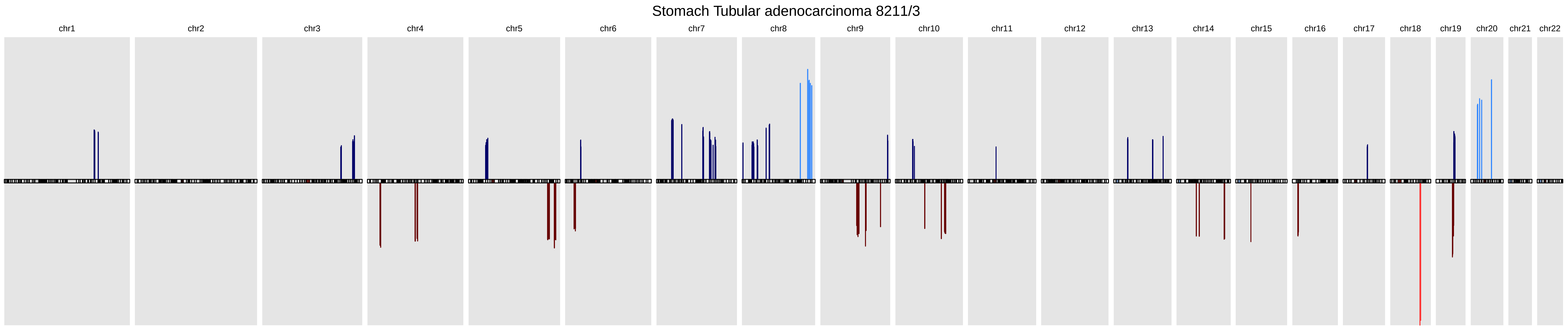

### sigGenes_full.pdf

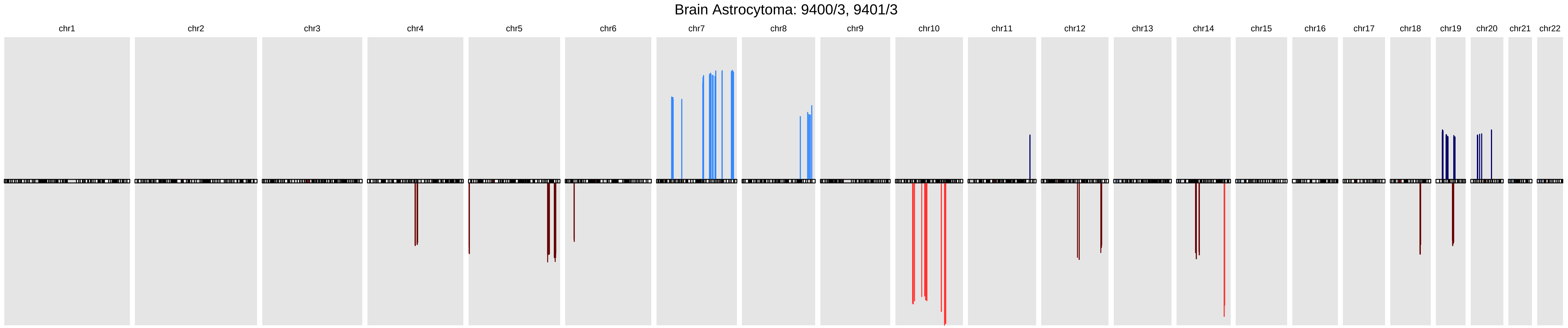

### sigGenes_full.pdf

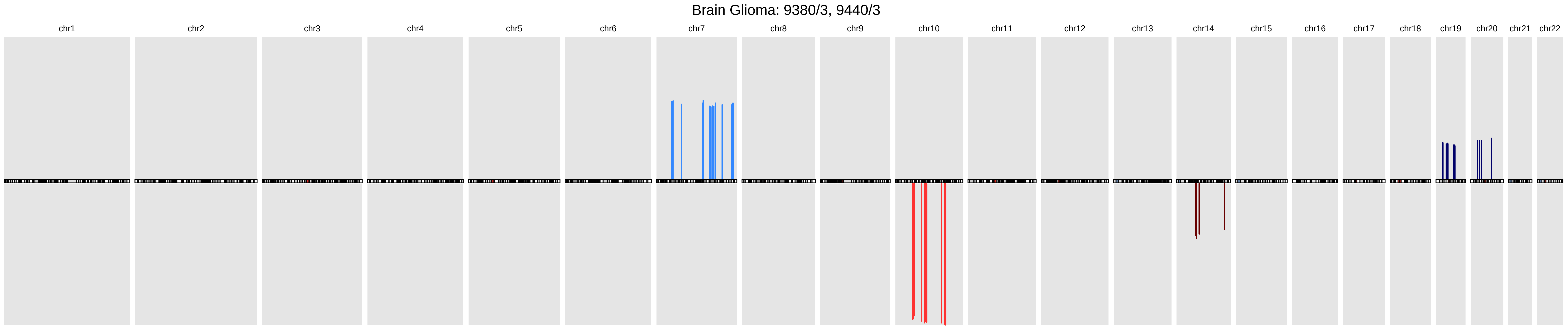

### sigGenes_full.pdf

#### Brain Oligodendroglioma: 9450/3, 9451/3

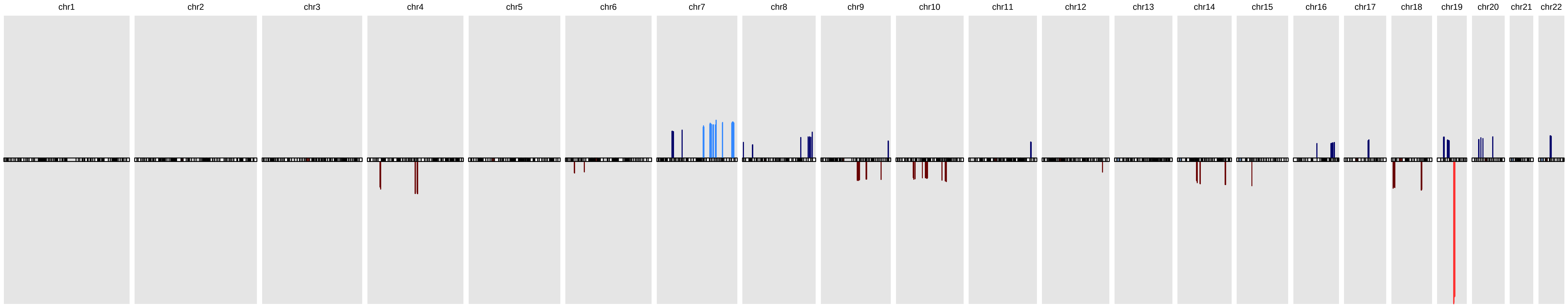

### sigGenes_full.pdf

#### Brain Mixed glioma: 9382/3

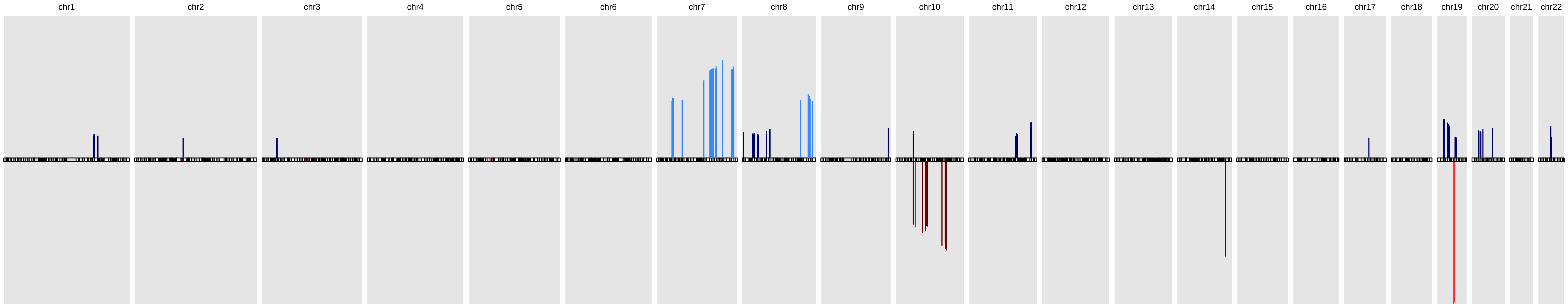

### sigGenes_full.pdf

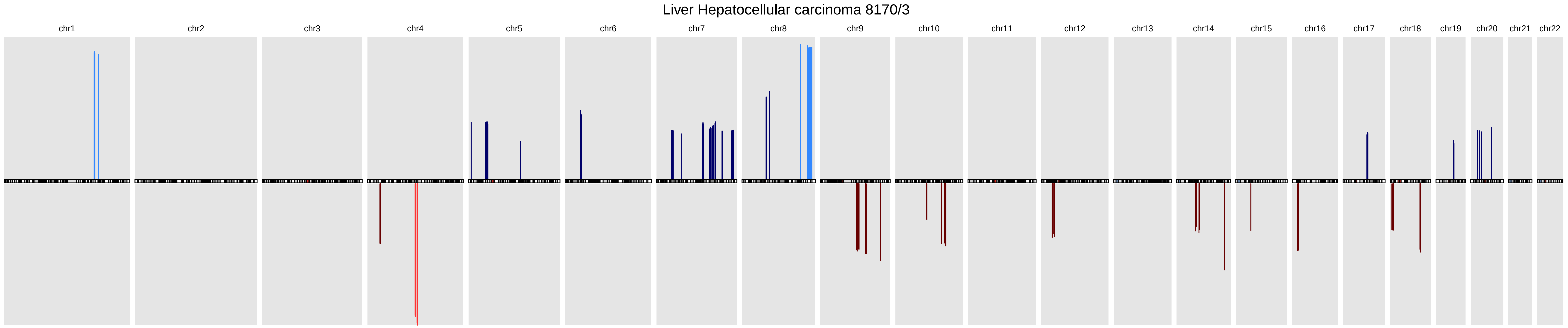

### sigGenes_full.pdf

#### Ovary Carcinoma: 8010/3, 8441/3, 8442/1

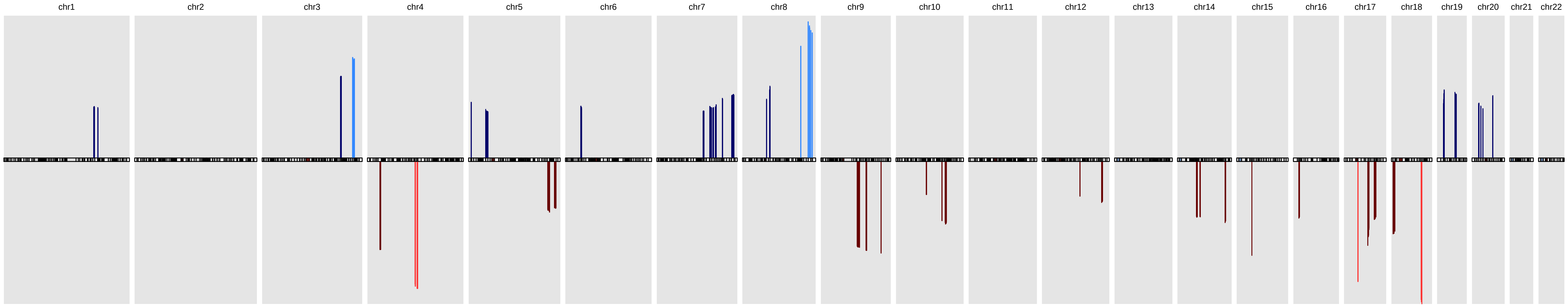

### sigGenes_full.pdf

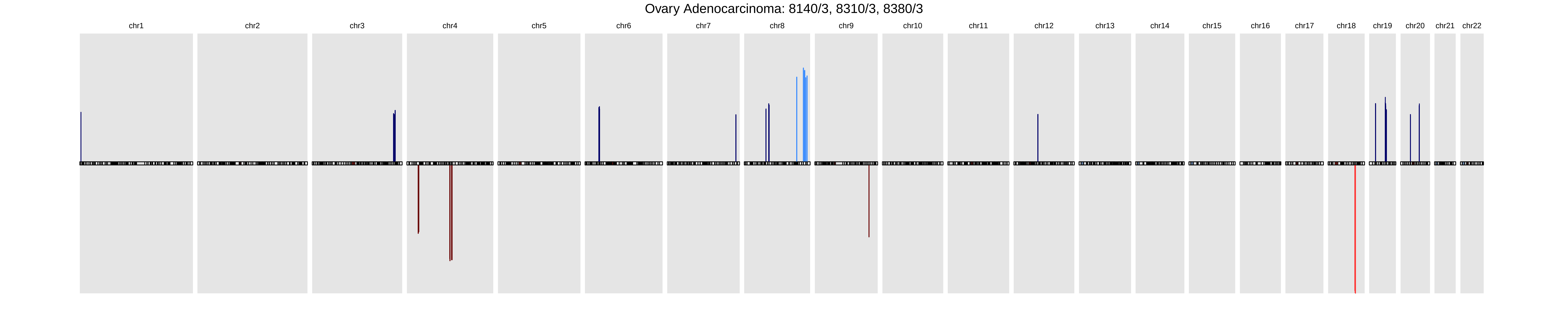

### sigGenes_full.pdf

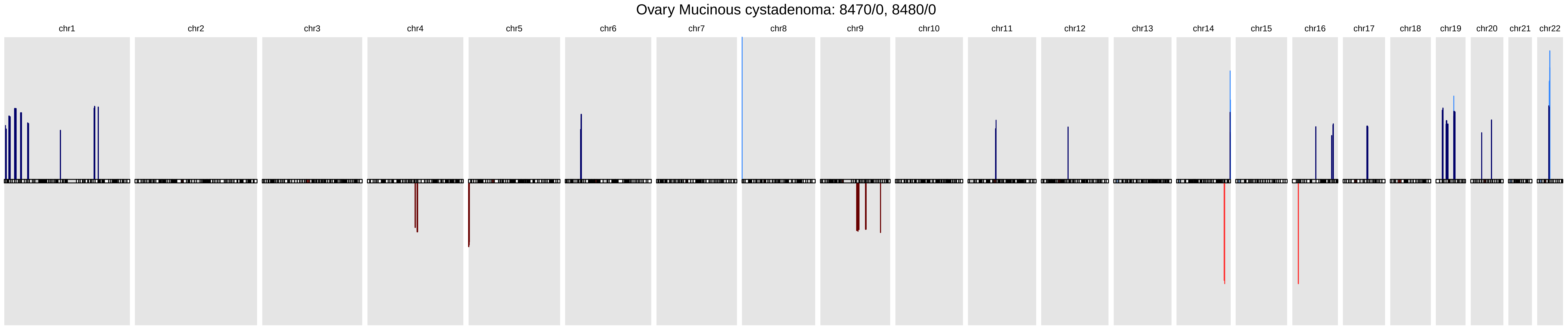
