## Supplementary Subtypes for "Signatures of Discriminative Copy Number Aberrations in 31 Cancer Subtypes"

Merged based on signatures, ■ is left out (less than 50).

| Tissue | Analyzed subtypes | Annotated subtypes | IDCO morphology | Samples |
|---|---|---|---|---|
| Brain | Glioma, malignant | Glioma, malignant | 9380/3 | 159 |
| | | Glioblastoma | 9440/3 | 1646 |
| | Astrocytoma | Astrocytoma | 9400/3 | 160 |
| | | Astrocytoma, anaplastic | 9401/3 | 179 |
| | Oligodendroglioma | Oligodendroglioma | 9450/3 | 221 |
| | | Oligodendroglioma, anaplastic | 9451/3 | 105 |
| | Primitive neuroectodermal tumor | | 9473/3 | 82 |
| | Mixed glioma (Anaplastic oligoastrocytoma, Oligoastrocytoma) | | 9382/3 | 162 |
| | | Ependymoma | 9391/3 | 49 |
| Breast | Intraductal carcinoma, noninfiltrating, NOS | | 8500/2 | 82 |
| | Infiltrating duct carcinoma, NOS | | 8500/3 | 5657 |
| | Lobular carcinoma, NOS | | 8520/3 | 201 |
| | | Infiltrating duct and lobular carcinoma | 8522/3 | 28 |
| | | Infiltrating duct mixed with other types of carcinoma | 8523/3 | 14 |
| | | Inflammatory carcinoma | 8530/3 | 19 |
| | | Metaplastic carcinoma, NOS | 8575/3 | 16 |
| | | Pleomorphic carcinoma | 8022/3 | 14 |
| | | Mucinous adenocarcinoma | 8480/3 | 15 |
| | | Intraductal micropapillary carcinoma (8507/2) | 8507/3 | 43 |
| Cerebellum | Medulloblastoma, NOS | Medulloblastoma, NOS | 9470/3 | 1573 |
| | | Desmoplastic nodular medulloblastoma | 9471/3 | 72 |
| | | Large cell medulloblastoma | 9474/3 | 37 |
| Colon | Adenoma, NOS | | 8140/0 | 63 |
| | Adenocarcinoma, NOS | | 8140/3 | 1643 |
| | Adenocarcinoma, intestinal type | | 8144/3 | 53 |
| | Mucinous adenocarcinoma | | 8480/3 | 62 |
| Kidney | Clear cell adenocarcinoma, NOS | | 8310/3 | 930 |
| | Renal cell carcinoma, NOS | | 8312/3 | 323 |

| Tissue | Analyzed subtypes | Annotated subtypes | IDCO morphology | Samples |
|---|---|---|---|---|
| | | Renal cell carcinoma, chromophobe type | 8317/3 | 15 |
| | | Oxyphilic adenoma | 8290/0 | 17 |
| Liver | Hepatocellular carcinoma | | 8170/3 | 371 |
| Lung | Carcinoma, NOS | Carcinoma, NOS | 8010/3 | 84 |
| | | Large cell carcinoma, NOS | 8012/3 | 54 |
| | Adenocarcinoma | Adenocarcinoma | 8140/3 | 1103 |
| | | Adenocarcinoma with mixed subtypes | 8255/3 | 109 |
| | Small cell carcinoma, NOS | | 8041/3 | 155 |
| | Non-small cell carcinoma | | 8046/3 | 1725 |
| | Squamous cell carcinoma, NOS | | 8070/3 | 518 |
| | | Squamous cell carcinoma, uncertain | 8070/1 | 12 |
| | Bronchiolo-alveolar adenocarcinomas | Bronchiolo-alveolar adenocarcinoma, NOS | 8250/3 | 24 |
| | | Bronchiolo-alveolar carcinoma, non-mucinous | 8252/3 | 19 |
| | | Bronchiolo-alveolar carcinoma, mixed mucinous and non-mucinous | 8254/3 | 18 |
| | | Papillary adenocarcinoma, NOS | 8260/3 | 29 |
| | | Mucinous adenocarcinoma | 8480/3 | 14 |
| | | Acinar cell carcinoma | 8550/3 | 21 |
| | | Adenosquamous carcinoma | 8560/3 | 16 |
| Ovary | Carcinoma | Carcinoma | 8010/3 | 761 |
| | | Serous carcinoma | 8441/3 | 964 |
| | | Serous tumor | 8442/1 | 45 |
| | Mucinous cystadenoma | Mucinous cystadenoma | 8470/0 | 57 |
| | | Mucinous adenoma | 8480/0 | 60 |
| | Adenocarcinoma | | 8140/3 | 111 |
| | | Clear Cell Adenocarcinoma | 8310/3 | 24 |
| | | Endometrioid adenocarcinoma | 8380/3 | 11 |
| Prostate | Adenocarcinoma | | 8140/3 | 916 |
| | | Carcinoma, NOS | 8010/3 | 16 |
| | Malignant melanoma, NOS | Malignant melanoma, NOS | 8720/3 | 1030 |
| | | Nodular melanoma | 8721/3 | 22 |
| | | Amelanotic melanoma | 8730/3 | 16 |

| Tissue | Analyzed subtypes | Annotated subtypes | IDCO morphology | Samples |
| --- | --- | --- | --- | --- |
| Skin | | Pigmented dermatofibrosarcoma protuberans (Bednar tumor) | 8833/3 | 11 |
| | | Mycosis fungoides (Pagetoid reticulosis) | 9700/3 | 32 |
| | | Epidermoid carcinoma (Squamous cell carcinoma), NOS | 8070/3 | 18 |
| | | Keratinizing (Squamous cell carcinoma) | 8071/0 | 11 |
| Stomach | Adenocarcinoma | | 8140/3 | 763 |
| | Gastrointestinal stromal sarcoma | | 8936/3 | 175 |
| | Adenocarcinoma, intestinal type | | 8144/3 | 83 |
| | Carcinoma, diffuse type | | 8145/3 | 57 |
| | Tubular adenocarcinoma | | 8211/3 | 82 |
| | | Mucinous adenocarcinoma | 8480/3 | 21 |
| | | Signet ring cell carcinoma | 8490/3 | 16 |
| | | Carcinoma | 8010/3 | 19 |
| | | Adenoma, NOS | 8140/0 | 17 |
| | | Adenocarcinoma in situ, NOS | 8140/2 | 19 |
